# An Efficient and Principled Model to Jointly Learn the Agnostic and Multifactorial Effect in Large-Scale Biological Data

**DOI:** 10.1101/2024.04.12.589306

**Authors:** Zuolin Cheng, Songtao Wei, Yinxue Wang, Yizhi Wang, Q Richard Lu, Yue Wang, Guoqiang Yu

## Abstract

The rich information contained in biological data is often distorted by multiple interacting intrinsic or extrinsic factors. Modeling the effects of these factors is necessary to uncover the underlying true signals. However, this is challenging in real applications, because biological data usually consist of tens of thousands or millions of factors, and no reliable prior knowledge is available on how these factors exert the effect, to what degree the effect is, as well as how they interact with each other. Thus, the existing approaches rely on excessive simplification or unrealistic assumptions such as the probabilistic independence among factors. In this paper, we report the finding that after reformulating the data as a contingency tensor the problem can be well addressed by a fundamental machine learning principle, Maximum Entropy, with an extra effort of designing an efficient algorithm to solve the large-scale optimization problem. Based on the principle of maximum entropy, and by constraining the marginals of the contingency tensor using the observed values, our Conditional Multifactorial Contingency (CMC) model imposes minimum but essential assumptions about the multifactorial joint effects and leads to a conceptually simple distribution, which informs how these factors exert the effects and interact with each other. By replacing hard constraints with expected values, CMC avoids the NP-hard problem and results in a theoretically solvable convex problem. However, due to the large scale of variables and constraints, the standard convex solvers do not work. Exploring the special properties of the CMC model we developed an efficient iterative optimizer, which reduces the running time from infeasible to minutes or from days to seconds. We applied CMC to quite a few cutting-edge biological applications, including the detection of driving transcription factor, scRNA-seq normalization, cancer-associated gene identification, GO-term activity transformation, and quantification of single-cell-level similarity. CMC gained much better performance than other methods with respect to various evaluation criteria. Our source code of CMC as well as its example applications can be found at https://github.com/yu-lab-vt/CMC.

**One-Sentence Summary:** CMC jointly learns intertwined effects of numerous factors in biologival data and outperform existing methods in multiple cutting-edge biological applications.

## Main Text

The volume of biological data is rapidly growing with the fast advancement of techniques, containing rich information about biological-related quantities under various conditions (1–4). The observed biological data are often influenced by numerous intrinsic or extrinsic factors at the same time, which may distort or even bury the impact of variables of real interest. For instance, mutation data collection (Fig. 1A Application 1), which documents the DNA mutations occurring in each gene for each individual cancer patient, is widely used to study the relationship between mutation and cancer initiation or progression (*2, 5, 6*). However, the data is affected by intrinsic factors such as each patient’s genetic background, gene-related factors, and types of DNA mutations (such as C:G or A:T transversions, etc.) (*2, 7*). As another example, transcription factor (TF)-gene binding state data (Fig. 1A Application 3) (*3*), which measures whether a TF binds to a gene, are simultaneously affected by tons of gene characteristics as well as tons of TF characteristics.

**Fig. 1.**
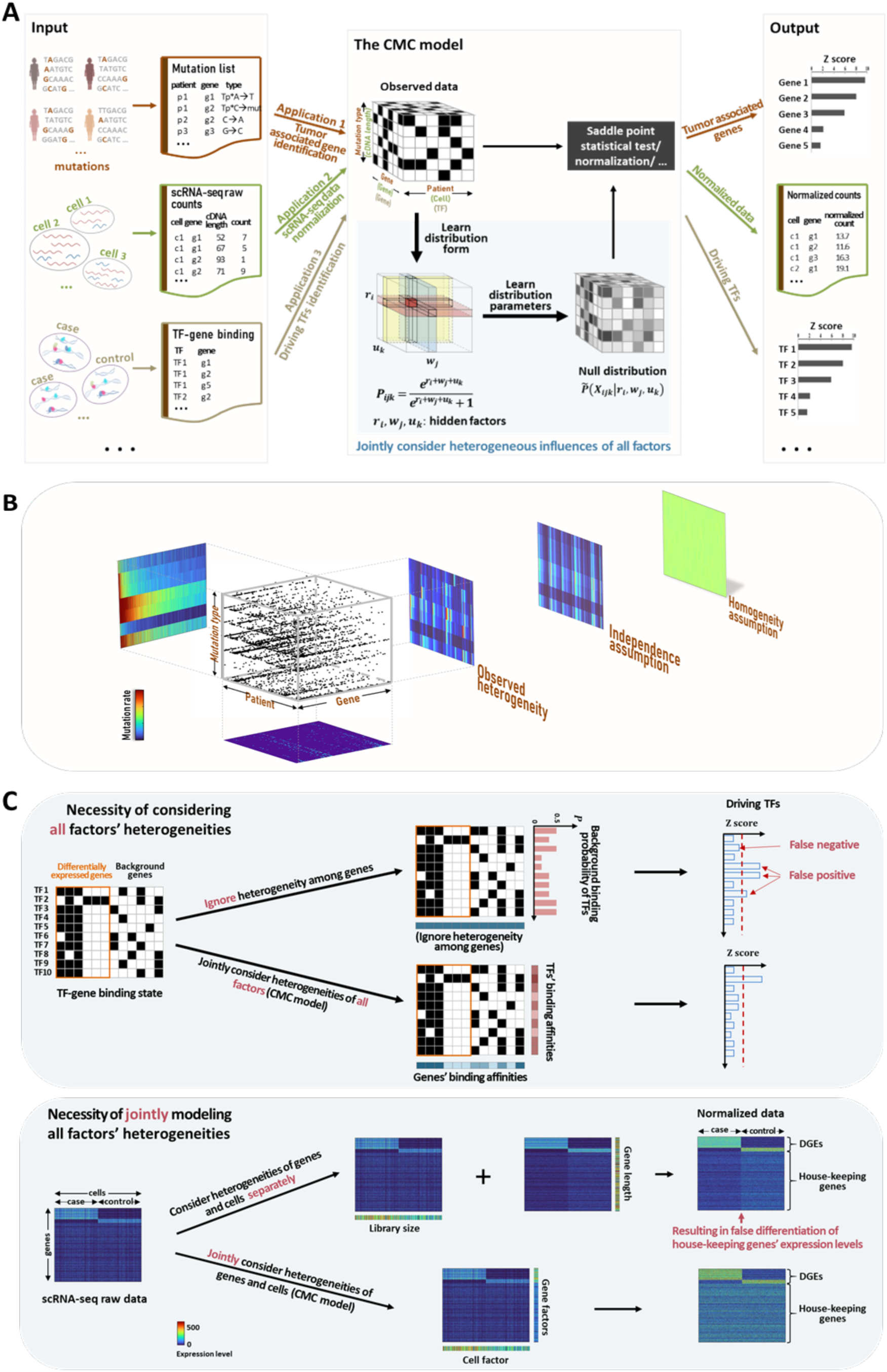
Introduction to the CMC model. (A) CMC model and its three example applications: Tumor associated gene identification (application 1); scRNA-seq data normalization (application 2); and driving TF identification (application 3). Left: for these applications, the values in their data (mutation list; scRNA-seq raw count; and TF-gene binding respectively) are often relevant to multiple factors (gene, patient, and mutation type factors in application 1; gene, cell, and cDNA length factors in application 2; and gene and TF factors in application 3) and are impacted by the individual-wise heterogeneity in each factor. Middle: the data of each application is reformulated to a multi-dimensional tensor in which each dimension corresponds to one factor class that exerts influence on the data values. For each entry in the tensor, the CMC model learns the optimal probability distribution form that reveals how the multiple factors jointly impact the value of this entry, as well as to what degree the effects are. The null distribution of each entry conditioned on the inferred impact strengthes is computed. Right: By comparing the observed data and its corresponding null distribution that incorporated with the heterogenous impact strengthes of all factors, CMC identifies the associated genes, normalizes out the unwanted variance, and detects the driving TFs for application 1∼3, respectively. (B) Heterogeneities among mutation data. Black dots in the 3D tensor indicate mutation events corresponding to specific gene/patient/mutation type. The heatmaps in three projected directions show the joint mutation rates of any two factors. Heatmaps of mutation rate under independence assumption and homogeneity assumption are also shown. (C) Necessity of jointly consider all factors’ heterogeneities. (Top) In the application of driving TFs identification, results are compared between jointly considering the heterogeneities of gene and TF factors and ignoring the heterogeneity of gene factor. Ignoring gene factor results in false positive and negatives. (Bottom) In the scRNA-seq normalization, results are shown for jointly or separately considering the heterogeneities of gene and cell factors. The separate application of normalization on different dimensions leads to problems in housekeeping genes

Explicitly modeling the effects of a large number of factors is challenging, as no reliable prior knowledge is available on how these factors exert their effects, as well as to what degree the effects are. What’s more, the numerous factors’ effects are often intertwined, and call for joint consideration of all the factors (Fig. 1B) (*2, 8, 9*). The necessity of jointly considering the heterogeneities in all factors is further confirmed by the consequences of either ignoring a part of the factors or considering the factors separately (Fig. 1C). Yet, disentangling multiple mutually correlated factors from data alone is not a trivial problem, as we are usually agnostic on how the factor interact with each other. As a consequence, existing approaches make oversimplified or unrealistic assumptions, either considering all factors in a given group to have the same effect or estimating each factor’s effect individually and combining multiple factors’ effects with an arbitrarily given formula (*2, 5, 8, 10–14*).

A case in point for this unfortunate scenario is the driving TF identification for a query gene set, an application that leans heavily on TF-gene binding state data. All of the state-of-the-art studies (*10–13*) treated the genes equally, likely due to the mathematical convenience. However, different genes obviously have different affinities of being bound by a TF. Another example is the normalization of scRNA-seq data. (*8, 9*) normalized out the gene factor and/or the cell factor independently, while (*15–18*) partially considered the gene factor when estimating the cell factor. The last example is cancer-associated gene identification which relies on mutation data. A formerly prevalent method, MutSig1.0 (*5*), ignored factors that exhibit gene-wise or patient-wise heterogeneous impacts and solely considered mutation types. An improved method, MutSigCV (*2*), acknowledged the importance of considering the different impacts of each factor, but treated these factors independently and used an unfounded multiplicative function to achieve a joint effect, which still resulted in many false-positive findings.

In this work, we show that this problem can be well addressed with a fundamental principle in the machine learning field, *maximum entropy*, by reformulating the data as a contingency tensor with each dimension corresponding to a factor class, and switching to a new type of constraint. Here, factor class refers to a group of factors that can be defined by users but should be exclusive to each other. For example, the cancer mutation data, which have three factor classes (patient, gene, and mutation type), can be reformulated as a high dimensional tensor (Fig. 1A). And naturally, within each factor class, a factor represents a particular individual, such a particular patient or gene.

We name our proposed model Conditional Multifactorial Contingency (CMC). Under the guidance of maximum entropy, CMC aims to learn the joint probability distribution of each entry in the contingency tensor with the expectations of the margins along each dimension fixed to the observed values. This is a large-scale optimization problem with potentially billions of variables and millions of constraints. By applying the Lagrangian method, we obtained an unconstrained optimization problem with a much-reduced number of variables. Interestingly, the impact strengths of factors can be well depicted by Lagrange multipliers, which naturally emerge during the optimization process.

However, to make the model practically useful, there are two issues to be resolved. 1) Efficiently solve this large-scale optimization problem. Even though the Lagrangian method has simplified the problem significantly, with tens of thousands of variables, solving it is still a daunting task. Indeed, we tried many existing solvers and all of them are impractical. By fully exploring the special structure of the CMC model, we developed a highly efficient iterative strategy to solve the optimization problem. Our strategy exhibits a remarkable speed and memory advantage, being more than 100,000 times faster than standard convex solvers and using 1,000 folds less memory when applied to real-world datasets. 2) Effectively conduct statistical analysis with the probability generated by the CMC model. Notice that each entry in the contingency table has a different probability due to factors’ heterogeneous impacts. Usually, we are interested in a given part of the entries and perform hypothesis testing for the observed values. Yet, the p-value does not follow an analytical form when probabilities are not equal. We resolved this issue by leveraging the saddle point theory and developing a numerical scheme for computing the p-value.

Here we present the CMC model with more details. Fig. 1A gives an overview of the CMC model. The CMC model reformulates the biological data into a multi-dimension contingency tensor in which each dimension corresponds to one factor class whose individual-wise heterogeneous effects contribute to the data’s variance. A probability distribution for each entry of the tensor is then learned. The distribution comes from joint consideration of all factor classes so as to reveals how these factor classes collectively impact the value of that entry. For statistical inference, this distribution can be used as the null distribution of the effect of heterogeneity. Instead of directly learning the distribution of each entry, CMC learns a hidden vector for each factor class, and each element of the vector measure how strong the effect of each individual (e.g., a gene, a patient, etc.) contributes to a value of interest: mutation count, binding state, or expression level. Therefore, each vector, with various values in its elements, models the heterogenous effects of the corresponding factor calss. The estimated effect strength of an individual can be regarded as an overall measure of this individual’s intrinsic characteristic w.r.t. the values of interest. For example, in the mutation count dataset, the estimated effect strength of a patient reflects his/her inherent vulnerability to any mutation due to genetic background and/or any other contributors.

The CMC model estimates the distribution based on the marginal totals in each dimension. A marginal total is the sum of all entries corresponding to one index (an individual or a factor) in one dimension. For each factor class, the marginal total reveals the impact strength of each factor to the observational event of interest, while the differential information of margins reflects the individual-wise heterogeneous effects within the factor class. Taking the mutation count tensor as an example, a larger marginal total of a patient/gene implies a higher vulnerability of this patient/gene to any DNA mutation.

With no extra assumption of the observed values other than the effect of all factors, the probability distribution of the entries in an 𝑁 × 𝑀 × 𝑄 tensor can be captured by maximizing the entropy under the marginal expectation constraints:

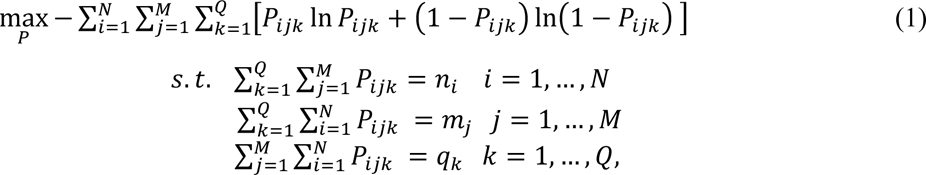

where 𝑃_𝑖𝑗𝑘_ is the probability of tensor entry (𝑖, 𝑗, 𝑘) to take value 1. We assume each entry follows the Bernoulli distribution. 𝑛*_i_*, 𝑚*_j_* and 𝑞*_k_* are the marginal totals of the 𝑖^th^, 𝑗^th^ and 𝑘^th^individual in the three dimensions, respectively. The marginal expectation constraints impose considerations for the heterogeneities, and the maximum entropy principle ensures no extra information is introduced. The CMC model is applicable on the tensor of any dimension. For simplicity, here we only present the three-dimensional case.

An alternative way to understand the role of the maximum entropy principle here is to search for the probability distribution of the tensor under the marginal expectation constraints and the consistency principle. The consistency principle is based on the concept that the solutions to a problem should be consistent if this problem can be solved in more than one way. Maximum entropy is a way to search for this unique distribution (*19*).

To solve Eq. 1, Lagrange multipliers {𝑟*_i_*}, {𝑤*_j_*} and {𝑞*_k_*} are introduced:

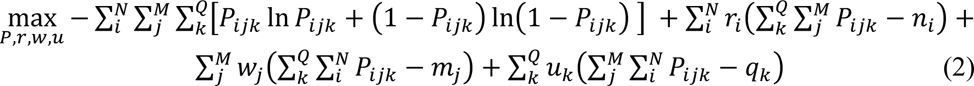

The optimal solution of Eq. 2 takes the form:

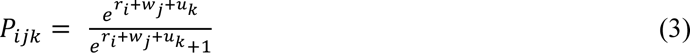

Eq. 3 reveals how the probability distribution is determined by the impact strengthes of all factors. It shows that the probability distribution of entry (*i, j, k*) is determined and only determined by multipliers with the *i*^th^ individual of the first factor, 𝑗^th^of the second, and 𝑘^th^of the last. This is consistent with our intuition that value in each entry should only be related to its corresponding individuals’ impact strengthes rather than others’. For example, the probability of a mutation occurring in a gene of a patient should only be related to this specific patient’s and this gene’s vulnerabilities rather than those of other patients or genes. Considering that 𝑃*_ijk_* is monotonically increasing with 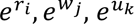 and that 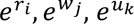 remain positive, 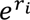 can be safely interpreted as a quantitative measure of the impact strengthes due to the intrinsic characteristic of individual (𝑖) within the first factor. The same applies to 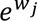 and 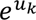.

The distribution form in Eq. 3 is consistent with the approximate mean of Fisher’s noncentral hypergeometric distribution of multiple binary variables. This distribution is a generalization of the hypergeometric distribution by allowing the modification of sampling probabilities with a weight factor. Indeed, the ratio of two individuals’ susceptivities within a factor corresponds to the ratio of these two individuals’ weights. However, Fisher’s noncentral hypergeometric distribution can only handle one factor, and even for the single factor, the weights of all individuals are required to be provided by users. Instead, in the CMC model, such susceptivities are automatically learned from the data itself.

Having obtained the probability distribution form (Eq. 3), the impact strengthes, or the parameters {𝑟*_i_*}, {𝑤*_j_*} and {𝑢*_k_*}, embedding in any given data can be learned through maximum likelihood estimation:

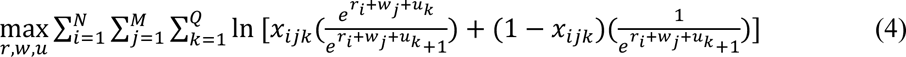

where 𝑥*_ijk_* ∈ {0,1} is the observed value in entry (𝑖, 𝑗, 𝑘) of the tensor.

According to Eq. 3, given the information of margins, the solution {𝑃*_ijk_*} is unique, as for any entry (𝑖, 𝑗, 𝑘), its 𝑟*_i_* + 𝑤*_j_* + 𝑢*_k_* is fixed (see Supplementary). The problem of searching 𝑁 × 𝑀 × 𝑄 variables {𝑃*_ijk_*} in Eq. 1 now converts into searching 𝑁 + 𝑀 + 𝑄 variables {𝑟*_i_*}, {𝑤*_j_*} and {𝑢*_k_*} in Eq. 4, which simplified the problem significantly. However, due to the large scale of variables and constraints, searching the global solution is still infeasible in many real applications. By fully exploring the special structure of the CMC model, we proposed a highly efficient iterative strategy (Fig. 2A, Supplementary Algorithm 1) and initialization strategies (see Supplementary) to solve the optimization problem. The former strategy aims to update variable sets {𝑟*_i_*}, {𝑤*_j_*} and {𝑢*_k_*} are updated respectively, based on an observation that: dependency among variables within a variable sets (say {𝑟*_i_*}) disappears, if the other variable sets are fixed. In other words, variables, 𝑟_1_, …, 𝑟*_N_*, can be independently optimized. And for any specific variable, 𝑟*_i_* for example, the solution can be obtained by Newton’s method. The latter strategies aim to get a good initialization for Newton’s method when solving a specific variable. They eighter utilize solution from history or from other variables belonging to the same variable set. Borrowing information from other variables is feasible thanks to the careful design of the algorithm after fully exploring the special structure of CMC model (see Supplementary). Our algorithm is guaranteed to converge to the global optimal solution (see Supplementary), and exhibits a remarkable speed and memory advantage, being more than 100,000 times faster than standard convex solvers and using 1,000 folds less memory when applied to real-world datasets (Fig. 2D).

**Fig. 2.**
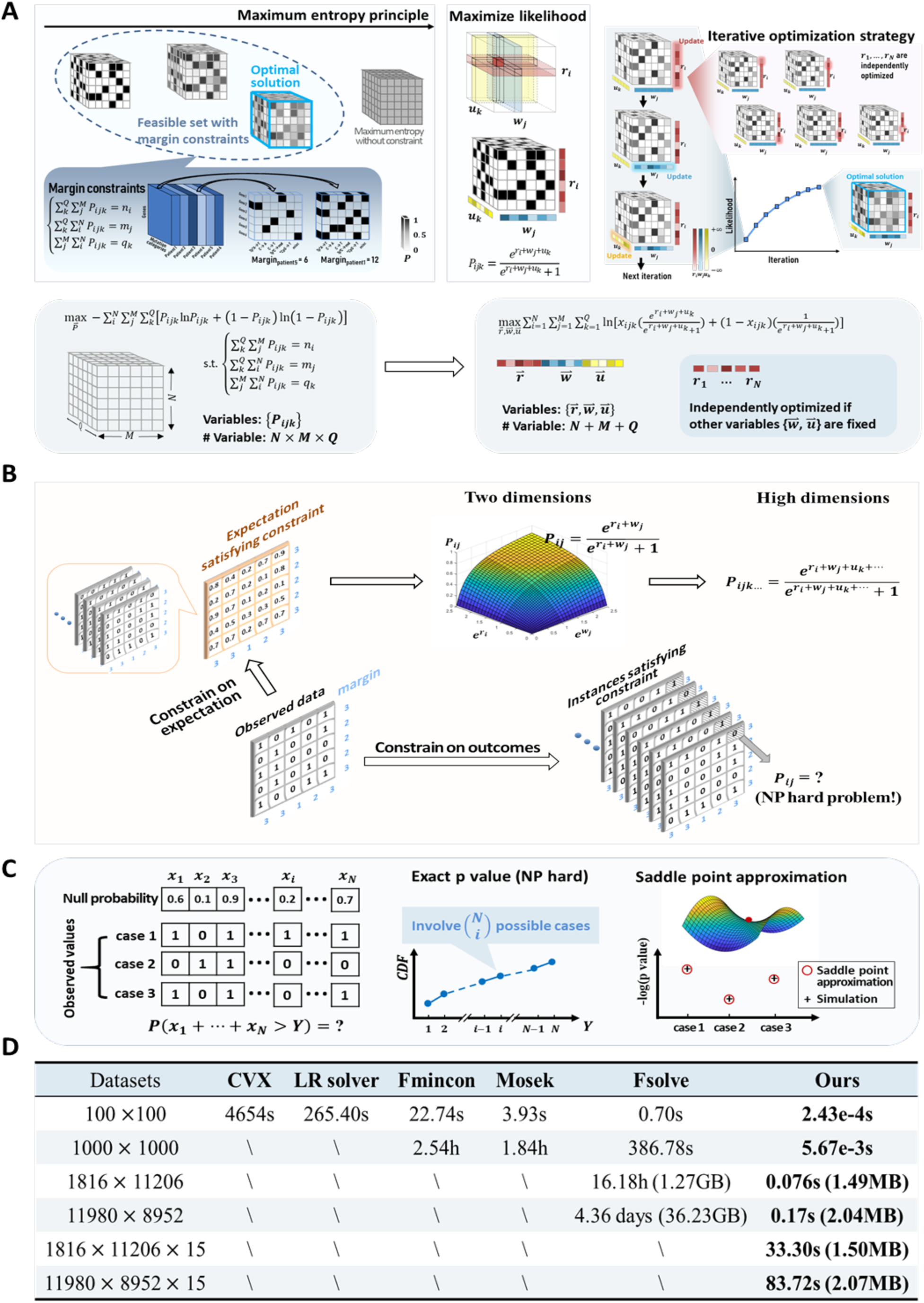
The CMC model. (A) Framework of CMC model. We took the 3D mutation count tensor as an example. A larger marginal total of a patient implies a higher vulnerability of this patient to the DNA mutations. With no extra assumption of the observational event of interest other than the heterogeneities in all factors, the heterogeneity induced null probability distribution of the tensor’s entries can be captured by maximizing the entropy under the marginal expectation constraints. Having obtained the form of probability distribution, the “susceptivity” of each individual in each factor is learned from the data through maximum likelihood estimation. The optimization problem can be quickly solved by iteratively optimizing variables corresponding to each dimension. During each iteration, variables of any specific dimension can be independently solved (in parallel) with variables of other dimensions fixed. (B) Constraints on expectation. CMC applies the marginal constraints on the sum of expectations of corresponding entries. The marginal constraints on the outcomes result in an NP-hard problem, while the constraint on expectation leads to a simple distribution form for any entries and can be easily extended to high dimensional tensors. (C) Saddle point approximation. Given the expected value of each entry in the tensor, applications like enrichment analysis need to check whether the observed values in a set of entries are significantly higher than the corresponding expected values. This procedure involves the statistical test on multiple entries wherein the expected values are typically different. Since computing the exact p-value is an NP-hard problem, saddle point approximation is utilized here to well approximate the exact *p*-value. (D) Efficiency comparison of different optimization solvers. In which, “LR solver” is short for logistic regression solver; “\” means either out of memory or running time is too long (> 5 days) to be recorded. The last four rows represent the data sizes coming from real studies.

One contribution and strength of this work lies in the type of marginal constraints that CMC employs. Instead of constraining the problem on the exact margins as noncentral hypergeometric distribution or other works did, CMC constrains the problem on the expectations of margins (Fig. 2B). The exact margin constraints result in an NP-hard problem, while the margin expectation constraints lead to a simple form of distribution (Eq. 3), which can be easily extended to high dimensional tensors.

The null distribution of the tensor can be extensively applied to downstream analyses. In many tasks, such as driving TF identification and cancer-associated gene detection, people are interested in querying whether some observations are significantly higher than the corresponding expected values. This procedure involves the statistical test on multiple entries jointly. The statistical test on entries sharing the same expected value is straightforward, with the null distribution being Binomial distribution. However, as CMC carefully takes care of the heterogeneities of all factors, the expected values of the entry set are typically different. In this case, the statistical test becomes extremely hard. For example, for a case with sum 𝑌 of the observed values in 𝑁 entries, there are 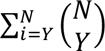 possible cases whose sums of observed values are larger than this specific case. And these 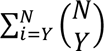 cases have different probabilities of being observed, making it an NP-hard problem to consider all these cases. The saddle point approximation is utilized here to well approximate the exact p-value (Fig. 2C; see Supplementary).

All the above discussion about the CMC model can easily be extended to high dimensional tensors, and tensors with values beyond binary numbers (such as probability values, and integer frequency count). It is also compatible with data with missing values (see Supplementary).

The CMC model is first applied to the single-cell RNA-seq (scRNA-seq) raw count data normalization. scRNA-seq is a sequencing technique used to measure the number of mRNA molecules of each gene in a single cell (*20*). To prepare the sequencing library, mRNA molecules of each cell are reverse transcripted to cDNA and then go through steps such as PCR amplification and fragmentation. The raw count is the number of reads (fragments) over a gene. During these processes, the sequencing output (raw counts) is impacted by artificial factors, including the cell factor (such as cell’s library size that will directly impact the total counts of the cell) and the gene factor (such as GC content of the gene’s sequence that may impact the efficiency of PCR amplification). Besides, we found that, for full-length sequencing techniques, such as Smart-seq2 (*21,22*), the cDNA-length factor also has a non-negligible impact on the final raw counts (Fig. S 3). cDNA length is defined as the sequence length of the cDNA after its reverse transcription from the template mRNA. A longer sequence implies more fragments to be generated after the fragmentation procedure and consequently larger raw counts in the final step. Typically, the cDNA length is equal to the gene’s mRNA that it is reverse transcribed from.

However, incomplete reverse transcription may occur and result in truncated cDNAs that are shorter than the typical one and will generate fewer fragments in the downstream steps (*23*). As a result, the corresponding mRNA tends to have smaller read counts. We observed such a non-negligible impact of the cDNA-length factor in scRNA-seq datasets from different labs (*24*, *25*).

To normalize the scRNA-seq raw counts, we constructed the raw counts into a tensor with three dimensions, corresponding to the gene factor, cell factor, and cDNA-length factor, respectively.

In the experiment, the cDNA length is inferred during the alignment process (see Supplementary). The CMC model was applied to jointly infer these three factors and then to normalize out the unwanted cell and cDNA-length factors. The gene factor is not normalized out from the data, because it includes information of gene’s average expression level across all cells. Users might be interested in such information.

More than 80% of values in a typical scRNA-seq data set are zero. However, a non-negligible part of them is not truly zero. Instead, due to the small number of RNA molecules in a single cell, many non-zero values cannot be detected (*26*). As there is no source information to distinguish truly zero from non-truly zero, we regarded all the zeros as missing values. The CMC model is compatible with data containing missing values (see Supplementary).

We use two criteria to evaluate the normalization performance: 1) the variance of each gene’s expression across a set of homogeneous cells; 2) the differentially expressed gene (DEG) detection results for two distinct cell types. The first criterion is a direct measure showing how much of the variance due to noise (both sequencing noise and biological noise) is removed after the normalization. The second one illustrates the overall impact of normalization on the downstream analysis like DEG detection; the biological signal in the dataset should not be falsely removed by the normalization.

The dataset used for this purpose is the scRNA-seq data of two cell types: immature microglia and proliferative-region-associated microglia (PAM), where the cell type labels are determined by clustering analysis based on cells’ expression profiles are also given (*24*). For the first criterion, Fig. 3A shows each gene’s variance change after normalization. Normalization using the CMC model reduces the variance of most genes to about half of their original variance. For comparison, Fig. 3B shows the variance changes after being normalized by a popular normalization method, total count (TC) (*5*), which considers the cell factor separately from the gene factor. It can be seen that, for the TC method, the variances reduce significantly for a small part of extremely highly expressed genes (such as “Rn45s” in this dataset). However, except for this small part of genes, the rest of the genes’ variances are only slightly changed. What’s more, variances of some ERCC spike-in controls, which are supposed to have small variances, even became larger after normalization by TC method. This is because the total count of each cell is dominated by a small number of extremely highly expressed genes. Normalizing out the cell factor without jointly considering such heterogeneity among genes will result in the normalization result favoring those highly expressed genes. We compared the variance change after normalizing by CMC and TC methods in Fig. 3C. As can be seen, compared with the TC method, CMC reduces the variance of most genes, except for a small part of genes. This small part of genes is mainly contributed by the genes with large 0 rates (typically corresponding to the genes with small detected values and suffer from large detection noise). Except that, a few other genes (e.g., genes with small 0 rates) also result in larger variances after normalization. We specifically inspect the genes on the left part (0 rates <0.15) in which many important genes are located for this dataset. It can be seen that, compared with the TC method, after normalization by the CMC model, most of the genes with larger variances are the differentially expressed genes (DEGs) reported by (*24*); meanwhile, the ERCC RNA spike-in controls have much smaller variance. It implies that the power of the CMC model in terms of reducing the variance of housekeeping genes while enlarging those of DEGs’. Fig. 3D summarizes the genes’ variance change after the normalization performed by the CMC model and other normalization methods, TC(*5*), TMM(*15*), RUV(*16*), UQ(*6*), SCnorm(*14*), Scran(*17*), and Census (*18*). It can be seen that the CMC model outperforms all the other methods in terms of reducing genes’ variances. As for the second criterion, we performed DEG analysis on the data normalized by different methods. We then compare the detected DEG (PAM vs. immature microglia) with the 937 “ground truth” DEGs. The ground truth DEGs downloaded from (*24*) are identified by comparing the expression of about 300 “pure” PAM cells with the immature microglia. The partial receiver operating characteristic (pROC) curve in Fig. 3E shows the CMC model significantly outperforms others on the basis of true positive rate again false positive rate. We also specifically examined the number of DEGs to be detected under the threshold of FDR<0.05. Results in Table S1 illustrate that the normalization by CMC model helps reveal more significant DEGs than all other normalization methods.

**Fig. 3.**
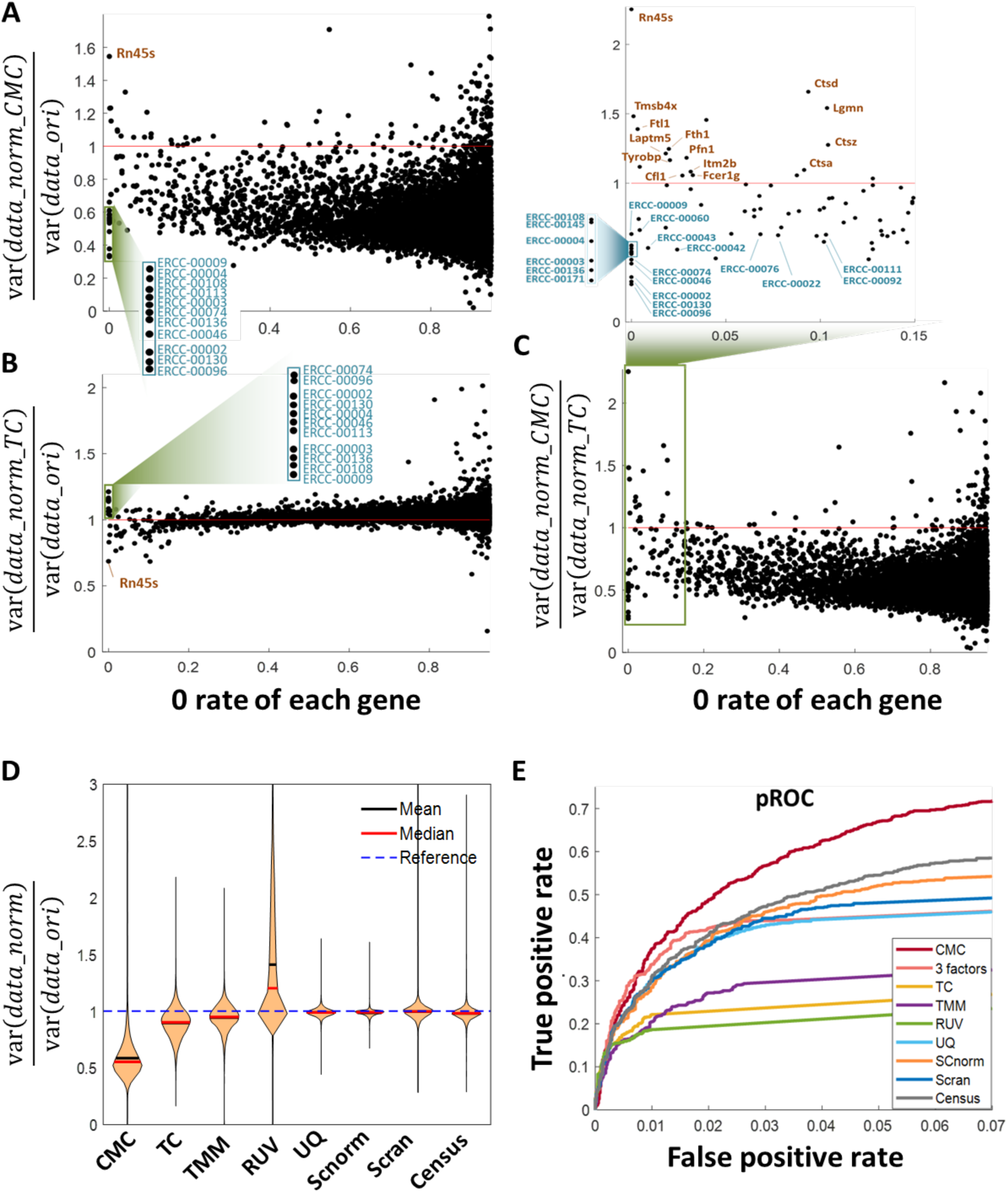
scRNA-seq data normalization. (A-B) Variance change of each gene across all cells before and after normalization (A: CMC model; B: Total count method). Each dot represents the variance change of an individual gene. The y-axis is the ratio of variance after normalization to that of before normalization. The variances are all computed on the log2-transformed normalized data. (C) Ratio of variance after normalization by CMC and TC mthods. The genes in the region of left side (0 rates < 0.15) are specifically inspected. Among them, the ERCC spike-in controls are legended in blue color texts; and for those genes with ratio larger than 1, we legended the DEGs in brown color texts. (D) Summary of all genes’ variance change after normalization via different methods. (E) Impact of normalization on downstream DEG analysis for two distinct cell types. The results are pROC curves of DEG detection after normalization via different methods.

Next, we applied the CMC model to enrichment analysis, including driving TF identification and tumor-associated gene identification.

The first one is to identify the driving TFs for a set of co-expression genes. TFs initiate and regulate the transcription of genes by binding to the enhancer or promoter sequences of their target genes. The basic idea behind the identification is to identify the TFs that bind to the co-expression genes more frequently than they would be at random. The key point of the identification is to accurately infer the expected by-random binding probability of each TF-gene pair. However, this is not trivial, as each TF or gene has different binding affinities (i.e., how likely a TF tends to bind any gene, or how likely a gene is bound by any TF). In addition, the TFs’ and genes’ binding affinities are cell-type specific (Fig. S 7). Previous driving TF identification methods (such as oPOSSUM 3.0 (*1*), Enrichr(*12*), BART(*11*), Lisa (*13*), etc.) often ignore the heterogeneity among genes’ binding affinities and treated all genes equally, which will falsely disregard the evidence from important genes and emphasize the importance of others. Here we applied the CMC model to jointly infer the cell-type-specific binding affinities of each gene and TF, and the random binding probability of any TF-gene pair conditioning on the TF’s and gene’s biding affinities. Fig. 4A shows the framework of the driving TF identification.

**Fig. 4.**
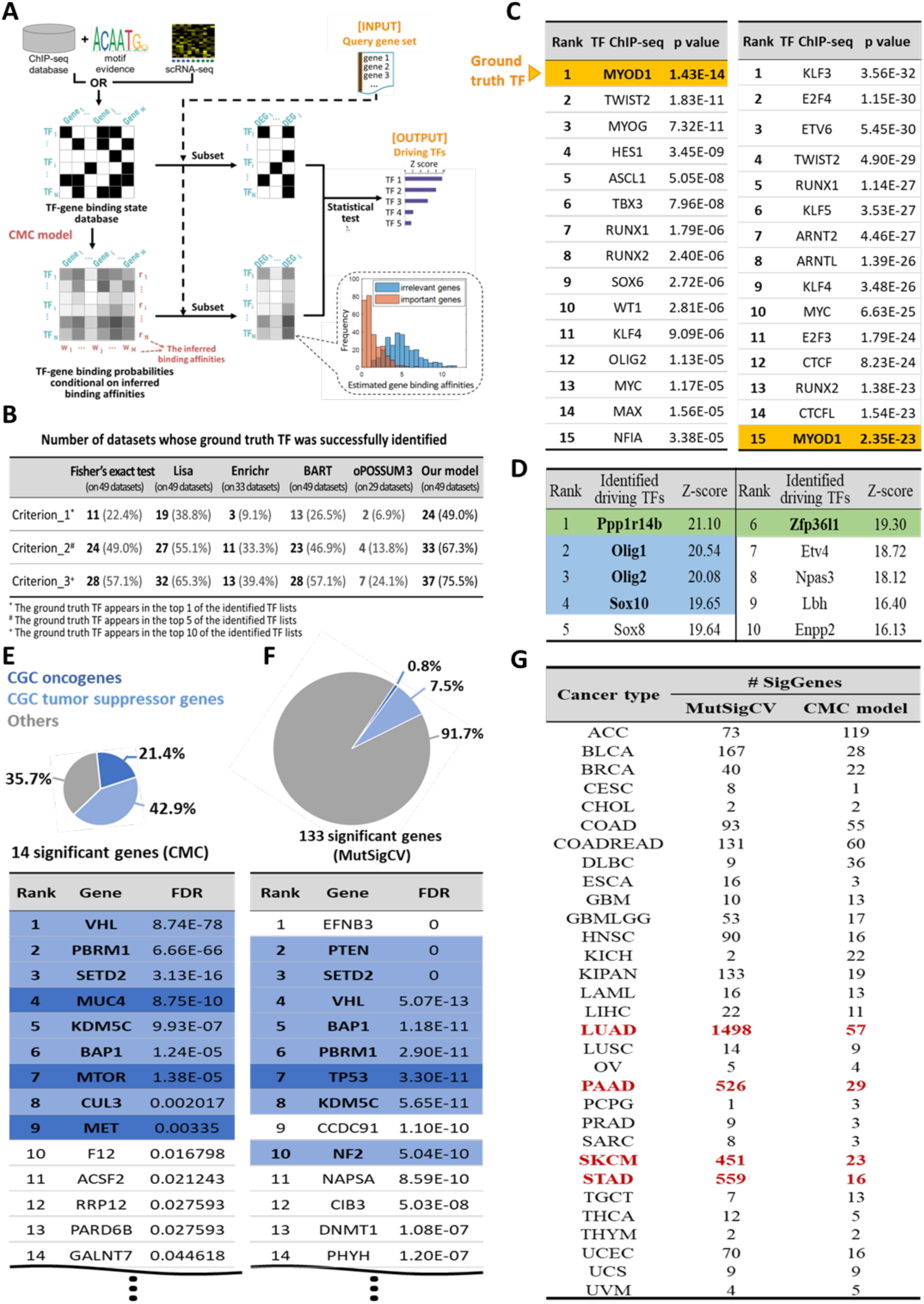
Driving TF and cancer-associated gene identification. (A-C) The framework and results of driving TF identification. (A) The framework. The input of this application is a co-expressed gene set and it outputs the identified driving TFs. The TF-gene binding state data is derived from ChIP-seq databases and stored in a 2D tensor. Each row/column corresponding to a TF/gene, respectively. The binding and non-binding states are distinguished by black and white colors. CMC model is applied to infer the binding affinities of each TF/gene, as well as the binding probability of each TF-gene pair. By comparing each TF’s observed binding state with the input co-expressed gene set and the corresponding binding probability, driving TFs are identified. (B) Summary of driving TF identification results on 49 benchmark datasets. (C) The significant TF identification results for an example benchmark dataset by the CMC model (Left) and the Fisher’s exact test (Right). The CMC successfully identified the ground truth TF, MYOD1, as the most significant driving TF. In the driving TF list identified by Fisher’s exact test, MYOD1 only ranks the 15^th^. It falsely identified some TFs whose ChIP-seq experiments are performed on embryonic fibroblasts or related cell types. Embryonic fibroblasts are the cell types that the input DEGs are detected in. (D) Top 10 identified TFs in a real study of identifying TFs that drive the oligodendrocyte fate and differentiation. Green color marked the well-known TFs; Blue color marked the two TFs that were confirmed by KO experiments. (E-G) Results of tumor-associated gene identification. (G) Number of significant genes identified for 31 tumor types by the CMC model and MutSigCV. mutSigCV identified more than 450 significant genes for LUAD, PAAD, SKCM, and STAD. (E-F) Tumor-associated gene identification results for KIPAN by CMC model (E) and mutSigCV (F). Among the 133 genes identified by mutSigCV, > 90% of them are neither CGC oncogenes nor tumor suppressor genes.

To measure the performance of the CMC model on driving TF detection, we used 49 benchmark datasets for testing (*13*). In each dataset, a single TF (or a TF family) is knocked out (KO) or overexpressed (OE) and the corresponding DEGs are identified. Therefore, we may input the DEGs into the identification model and check if the model can successfully identify the KO/OE TF that results in the differential expression of these genes.

Fig. 4B summarizes, among the 49 experiments, how many datasets whose ground truth TF is successfully identified. We used three criteria to define the “successful identification”, namely, the ground truth TF appears in the top 1, 5, or 10 of the identified TF lists. As shown in the Fig.s, by jointly considering the TF and gene factors, the CMC model surpasses all the other methods in all criteria. The second best is Lisa (*13*), which accounts for additional biological knowledge (i.e., genome-wide chromatin accessibility information).

We particularly checked the identification result in an example benchmark dataset with MYOD1 being knocked out and serving as the ground truth TF to be identified (Fig. 4C). Method based on Fisher’s exact test is a good comparison, as it used the same input as the CMC model but ignored the various impacts of genes’ binding affinities. As shown, the CMC successfully identified MYOD1 as the most significant driving TF, while in Fisher’s exact test method, it ranks 15^th^. We examined the binding affinities of important genes and irrelevant genes inferred by CMC model. Important gene refers to the muscle-specific genes from (27), as the target TF, MYOD1, binds to muscle-specific genes and promotes their expression (28). Irrelevant genes are referred to those frequently bound by many other TFs rather than the specific TF. As shown in Fig.4A, CMC method estimated a small binding affinity for these important genes, i.e., less possibility to be bound by random. Then in the downstream statistical test, these genes would contribute more to TF identification. Fisher’s exact test method, however, regards all important and irrelevant genes as equal, and consequently, fails to capture evidence from important genes while falling into the traps of irrelevant genes and TFs.

To further evaluate the performance of CMC model, we applied it to a real study case to identify the TFs that drive oligodendrocyte (OL) lineage. Identifying these TFs is vital for understanding the mechanism of glial progenitor differentiation and its related diseases, such as brain tumorigenesis. Fig. 4D shows the top 10 driving TFs identified by CMC model. In addition to the previously well-known TFs of OL fate commitment and differentiation, such as Olig1, Olig2, and Sox10 (ranked from 2^nd^ to 4^th^), the CMC models also identified other potential TFs, such as Ppp1r14b (ranked 1^st^) and Zfp36l1 (ranked 6^th^), whose roles in in regulating OL differentiation from NSC were confirmed via knock-out (KO) experiments (reported by (29)). Specifically, experiment showed that the number of OL lineage cells decreased significantly in mice with Zfp36l1 KO, while the number of other glial cells, astrocytes, increased dramatically, suggesting that Zfp36l1 controls the OL-astrocyte fate transition in the developing brain. Additionally, Zfp36l1 was found to play a key role in gliomagenesis. Similarly, the role of Ppp1r14b was also confirmed by Ppp1r14b-KO experiments. These results validate the effectiveness of our CMC-based driving TF detection model in real studies.

We also applied the CMC model for tumor-associated gene identification. Specifically, we want to identify the mutated genes responsible for the initiation and progression of a particular tumor. The basic idea is to detect the genes with significantly more mutations than expected by chance. We used the genomic mutation data from 8544 patients across all major tumor types from The Cancer Genome Atlas (TCGA) (*30,31*), in which the whole-genome sequencing is performed on the normal and tumor samples of each patient to identify the genomic mutations.

Again, estimating the expected mutations considering heterogeneity is critical. However, the mutation rates vary across patients, genome regions, and mutation categories (such as CpG transversions, C:G transitions, A:T transversions, etc.) Besides, such diversities between each factor (patient, gene, and mutation category) are twisted together (Fig. 1B). MutSigCV (*2*) reported the heterogeneities of these three factors. However, it treated the three factors independently and estimated the expected mutation number using the product of marginal relative rates of these factors. Here we applied the CMC model to jointly consider all these three factors. The framework of the detection is shown in Fig. S13. The mutation counts are stored in a tensor of three dimensions, corresponding to the patient, gene, and mutation category factors, respectively.

Fig. 4G summarizes the number of significant genes identified for 31 tumor types by the CMC model and MutSigCV. For tumor types LUAD, PAAD, SKCM, and STAD, MutSigCV identified more than 400 significant genes, while CMC resulted in a much small number of significant genes.

There are two reasons behind the abnormally large number of identified significant genes by MutSigCV. First, the “neighbor” genes MutSigCV identified were unreliable, resulting in inaccurately estimated background mutation rate of each gene. Specifically, MutSigCV directly estimated the background mutation rate for each gene (number of synonymous mutations and noncoding mutations divided by the sequence length). However, there is not enough data to confidently estimate the mutation rate for a single gene, especially for a short gene. Instead, MutSigCV had to estimate the average mutation rate for a specific gene and its “neighbors”. The neighbors were defined as genes that had similar characteristics, such as expression levels, DNA replication time, and open vs. closed chromatin status. What’s more, the gene characteristics are related to the cell types. The consequence of such a procedure is that MutSigCV’s results are quite sensitive to the user-provided gene characteristics. For example, using as input the overall gene characteristics (provided by the MutSigCV’s website), instead of cell-type-specific gene characteristics, the number of detected genes for “LUAD” changes dramatically from 1498 to 22. The CMC model learns the gene factor from the mutation-count tensor with more than 8000 patients, even these patients have different types of tumors as other factors, including patient factor, are jointly considered during the inference of gene factor. It does not need to identify each gene’s “neighbors”. The second reason is that MutSigCV considered the gene factors and patient factors independently, and the estimated mutation rates of each gene would be heavily impacted by some extreme patients. Specifically, for tumor “PAAD”, MutSigCV always identified about 400 significant genes regardless of the input gene characteristics. This is because, in “PAAD”, there is one patient who has extremely large mutation numbers (7794 nonsilent mutations and 5332 silent/noncoding mutations), while the mean nonsilent and silent/noncoding mutation number of the other 183 patients is 55.5 and 78.8 respectively. In MutSigCV, the estimation of each gene’s mutation rate was dominated by this specific patient. The CMC model, on the other hand, jointly considers the gene factor and patient factor. Therefore, the estimated gene factor will not be impacted by patients with extremely large mutations, as these patients would be assigned very large patient susceptivity values. The CMC model identified 29 significant genes for “PAAD”.

We particularly examined the identification results for “KIPAN” (Fig. 4E-F), where 14 and 133 genes are identified by CMC and MutSigCV. Among the identified genes by CMC, 21.4% of them are the CGC oncogenes (*32*) and 42.9% of them are the CGC tumor suppressor genes, while there are only 0.8% and 7.5% respectively for MutSigCV.

The forth application about the quantification of cell-to-group similarity. As scRNA-seq data is wildly used to exploring cell-to-cell heterogeneity and identifying new cell types (*24*, *33–41*), many studies struggled to answer a few common essential questions: 1) What is the relationship between this newly found cell type and a known cell type (*24,34–39*)? 2) Does a cell type newly found under one condition (e.g. disease model, species, tissue) also exist under another condition (*33, 36, 40, 41*)? The challenge here is a consequence of lacking well-accepted objective measurement of similarity between cell groups, especially at single-cell resolution and handling datasets generated from different sequencing platforms. In response to this, we propose a new quantitative metric that measures the similarity of a cell to a given target cell type w.r.t gene expression levels, as long as we have a set of signatures or DEGs (list of names) of this cell type. Such a score, named TySim, can provide a cell-to-group similarity assessment (single-cell-resolution) and hence accumulatively a group-to-group similarity assessment. TySim defines “a cell is similar to a cell type” if known signatures of the target cell type are significantly highly differentially expressed in the cell of interest. However, assessing the extent of highly differential expression for a gene in a cell is a non-trivial task, as it must be considered in the context of other genes. Furthermore, variations in sequencing depth can possibly make different cells’ sequencing counts incomparable; while different genes may enjoy different preferences in terms of sequencing, making genes incomparable in terms of measured gene expression level.

TySim employed the CMC to construct the distribution of each gene’s expression level in each cell if it is not differentially expressed, conditional on both the inferred sequencing affinity of a gene and that of a cell (a summary of the impacts of all possible gene/cell-related factors). From the null distribution given by CMC, a p-value is calculated for each cell summarizing the whole signature set, whose transformed form becomes our final similarity score. As a technical addition, a special design is introduced into the calculation of the TySim score upon consideration of the non-negligible dropout issue (*26*) in scRNA-seq data. Fig. 5A shows the framework of CMC-based TySim.

**Fig. 5.**
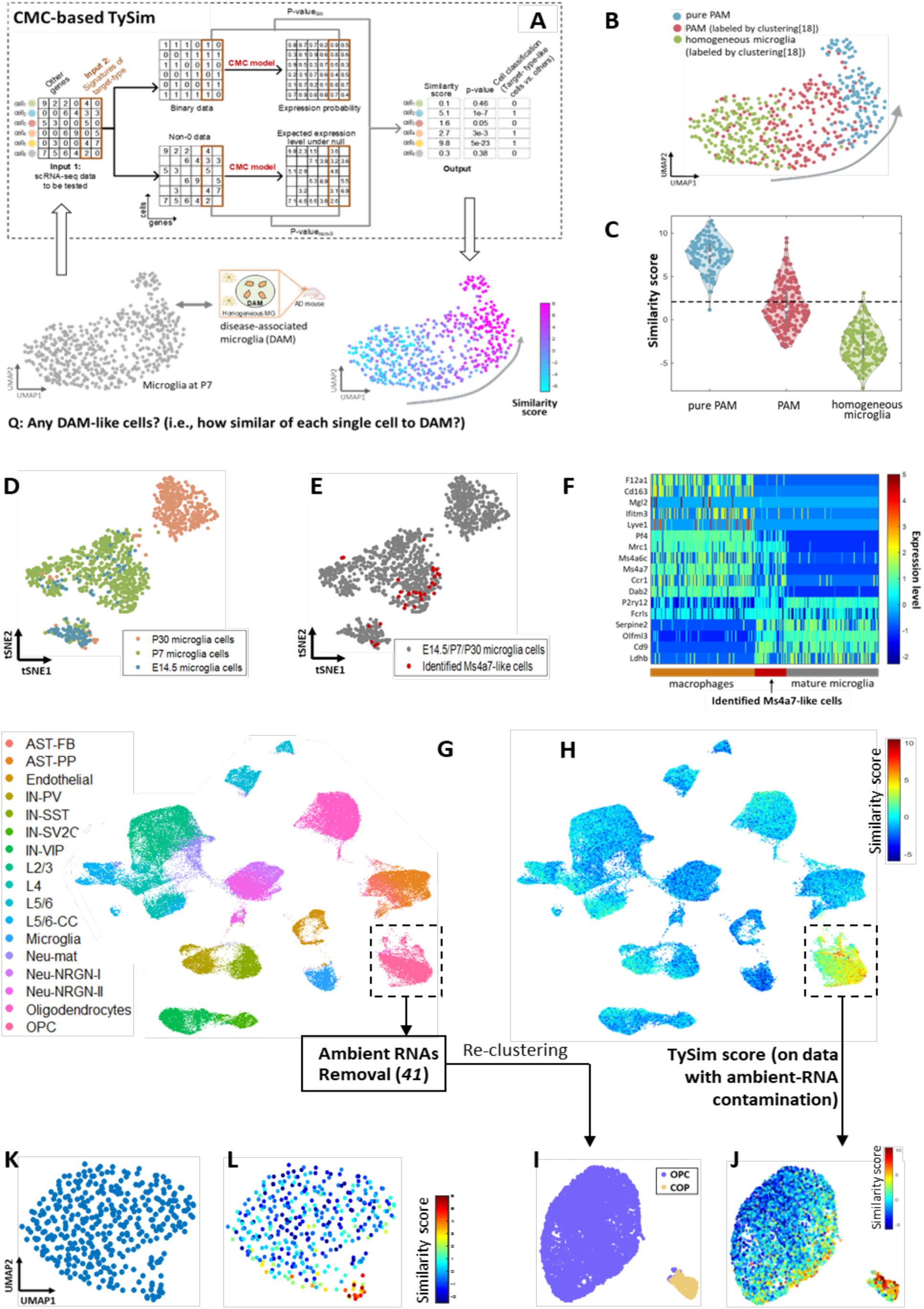
Framework of quantification of single-cell-level similarity and its experiment results in three study cases. (A) The framework. The model takes the scRNA-seq dataset to be tested and the signatures or DEGs of the target cell type as inputs, and outputs the similarity scores that indicate how similar of each cell to the target cell type. The inputted scRNA-seq dataset is separated into the binary part and the non-zero part to take care of the “dropout” phenomenon. For each of these two parts, we estimate the expected values of each item in the matrix under the null hypothesis. Each cell has two pvalues from the two independent hypothesis tests in binary data and in non-missing data. These two p-values are combined into a single p-value using Fisher’s method. We also provided z score to quantitative the similarity level. (B-C) First study case: TySim similarity score between microglia to DAM cell type. (B) UMAP plot showing cells to be tested, including the P7 microglia and GPNMB+CLEC7A+ cells (“pure PAM”). P7 microglia were clustered into PAM and homogeneous microglia (clustering labeled obtained from (*24*)). (C) TySim score distributions in pure-PAM, clustered-PAM, and clustered-homogeneous microglia groups, respectively. Beside, the bottom-right UMAP plot in (A) showing transition pattern in term of similarity to DAM. Each individual cell is color-coded by its TySim similarity score. (D-F) Second study case: TySim score reveals cell type neglected by clustering. (D) tSNE plot showing the 1236 microglia cells to be tested (*24*), including cells from three different ages: E14.5, P7 and P30. (E) The identified Ms4a7-like cells. (F) Heatmap of DEG expression of macrophages, Ms4a7-like cells (found by significant TySim similarity to Ms4a7+ microglia), and mature microglia. (G-L) Third study case: TySim Score Enables the Discovery of COP in data with contamination. (G) Human scRNA-seq dataset [89] to be tested. Cells are color-coded according to the cell types identified by (*43*). The data were contaminated by ambient RNAs and committed oligodendrocyte precursor (COP) was not found (*43*). (H) TySim scores that measures the similarity of each single cell to COP. (I) COP clustering identified by (*44*) after removing the ambient RNA contamination. (J) The same UMAP plot to show the correspondence between COP clustering by (*44*) and TySim similarity score. Here, cells are color-coded by the TySim similarity score of a cell to COP. Note that the similarity score quantification was performed on the original scRNA-seq dataset with ambient RNA contamination. (K-L) TySim similarity score of a cell to COP for another scRNA-seq data (*45*). This data are also contaminated by ambient RNAs. (*44*) fails to identify COP from this data even after remove the contamination (K). TySim score of each single cell to COP (B). Again, the scores were compute on the original data with contamination. The bottom-right cells with high scores are likely to be COP by inspecting the expression pattern of COP markers.

TySim is powerful in a few aspects. Firstly, to the best of our knowledge, it is the first quantitative metric of similarity towards a target cell type. Secondly, instead of traditional coarse group-to-group comparison, TySim relates each single cell to a given cell type, in which sense it is a “single-cell-resolution” metric and provides much finer information. Working on single cells also means that TySim does not require clustering and are free of the headaches in clustering, e.g. tiny cell group or inseparable groups. Lastly, since for the target cell type we only need a DEG name list that is no longer relying on any dataset, TySim has no troubles like an incompatibility between datasets from different sequencing platforms and the challenges in batch effect removal which become extremely challenging when datasets come from different sequencing platforms. This empowers TySim to work impressively well on cross-platform problems.

TySim similarity score can serve as an excellent facilitator or even enabler for diverse biological studies. Here, we present three example study cases using published real datasets, which also validate the effectiveness of TySim. In the first study (Fig. 5A-C), TySim score is confirmed to be able to quantitative the single-cell-to-target-cell-type similarity as expected, with expectation coming from prior knowledge about the cells and the cell types. Proliferative-region-associated microglia (PAM) is a subset of microglia mainly found in developing corpus callosum and cerebellar white matter (*24*). Reference (*24*) showed that PAM shares important signatures with degenerative disease-associated microglia (DAM) found in aging mouse (*33*). We tested TySim score using the scRNA-seq data published with (*24*) (Fig.5 B). The dataset contains homogeneous microglia, PAM, and “pure” PAM. We calculated TySim score of each cell in the dataset w.r.t. DAM cell type. The DEG list of DAM was obtained from (*33*). The expectation is that PAM cells should be much more similar to DAM than homogeneous microglia cells are.

Fig. 2A presents cell clusters found by (*24*) and the distribution of TySim similarity w.r.t. DAM in each of these cell clusters: “pure PAM”, PAM, and homogeneous microglia. TySim well distinguishes pure PAM and homogeneous microglia with almost no overlap in distribution (Fig. 5C). PAM labeled by clustering has a TySim distribution close to that of pure PAM with a small shift towards homogeneous microglia. This is expected since gene expression clustering try to assign discrete labels to every cell, including those in continuous transition state. Therefore, the PAM labeled by clustering may contain some cells in transition state. Overall, this result highlights the good power of TySim score to differentiate conceptually similar cell type (PAM) from others (homogeneous microglia). TySim similarity is further visualized at a single-cell resolution in Fig. 5A bottom-right. As each dot in the UMAP plot represents a cell, it is color-coded by its TySim w.r.t. DAM. We can see a clear smooth and almost monotonic spectrum of similarity from pure PAM to the furthest cells from pure PAM in the whole gene expression level space. This result demonstrates that TySim score successfully captures the order of true similarity, suggesting TySim’s capacity in illustrating the cell-type transition state at single cell resolution.

In the second study, for a microglia subset that plain clustering analysis failed to find due to very small amount of cells in this subset compared with the homogeneous microglia population, TySim accurately identifies the relevant cells (Fig. 5D-F). In this study, reference (*36*) questioned the counterintuitive absence of Ms4a7-expressing microglia subset (*37*) in the scRNA-seq data of (*24*). One hypothetical root cause is that clustering analysis is prone to overlook tiny cell groups, especially if they have relatively subtle differences from a much larger group(s) compared with the overall variations in the dataset. To verify this hypothesis, we applied the TySim score to the dataset of (*24*) along with the DEG list of Ms4a7-expressing microglia (Ms4a7+ microglia) given by (*37*). We found 32 cells that were statistically significantly similar to Ms4a7+ microglia (FDR < 0.05) (Fig. 5E). We then checked their characteristics to see if they are the same type. According to (*37*), Ms4a7^+^ microglia is an intermediate-state cell type between macrophage and mature microglia, and express part of markers of these two. We checked the expression pattern of the identified Ms4a7-like cells from (*24*)’s dataset. Fig. 5F shows that some makers of mature microglia are also highly expressed in Ms4a7-like cells, and so are some makers of macrophages, suggesting an intermediate state. This validates that Ms4a7-like cells found in the (*24*)’s dataset using TySim similarity well match Ms4a7^+^ microglia discovered in (*37*). It is worth noting that the amount of Ms4a7^+^ microglia is noticeably smaller in (*24*)’s dataset (30 out of 1236 microglia cells) than that in (*37*)’s dataset (more than 5,000 cells out of a total of more than 76,000 cells). The clustering technique was a valid method to discover this subset in the latter but failed in the former, probably due to small group size and subtle differences from others. In contrast, the TySim similarity score has a much stronger ability to reveal such small cell groups characterized by relatively subtle differences.

The third example study demonstrates that TySim enables the identification of cell type that was not discovered due to contamination in scRNA-seq data (Fig. G-L). In this study, we attempted to re-discover committed oligodendrocyte precursor (COP), a subset of oligodendrocyte, in human data, using TySim with a COP DEG list from mouse data. The DEG list of COP was got from (*42*) which studied the mouse oligodendrocyte lineage; while scRNA-seq data of human oligodendrocyte lineage was obtained from (*43*). Reference (*43*) failed to identify COP group in its dataset (Fig. 5G). We applied TySim to this dataset to calculate each cell’s similarity (Fig. 5H), which identified 170 cells that significant similarity to mouse COP. For validation purposes, we compared our result with that of a recently published study (*44*). Reference (*44*) managed to separate COPs from the larger population of oligodendrocyte precursor cells (OPCs) by cleaning up ambient RNAs in the scRNA-seq data before clustering analysis, and successfully identify the COP group in (*43*) data (Fig. 5I). Fig. 5I-J visualizes the correspondence between the two methods’ results by using the same UMAP dimensionality reduction. Fig. 5I color-codes the clusters given by (*44*), while Fig. 5J color-codes the TySim similarity score of each cell. It can be clearly seen that the COPs have significantly higher TySim scores than OPCs, meaning that TySim similarity successfully distinguishes COP. Besides, there’s a trend of similarity change in the OPC population, which is consistent with our knowledge about status transitions in the oligodendrocyte lineage. It is worth noting that our similarity evaluation was performed on the raw (*43*) dataset, without ambient RNA removal. This demonstrates a strong potential of TySim similarity score to work on various datasets even if there might be unknown contaminations inside.

The last application is the GO-term activity transformation. It’s widely acknowledged that scRNA-seq data are highly noisy and suffered from the drop-out effect, which limits its power in cell heterogeneity analyses. Compare with the expression value of every single gene in a cell, the overall expression pattern of a functional-related gene set is more reliable. Therefore, we propose a GO term activity transformation model to transform the scRNA-seq dataset into GO term activity score (Fig. 6A). Compare with the scRNA-seq dataset, which represents the expression level of each single gene in each cell, the GO term activity score matrix represents the activity level of each GO term (i.e., biological process) in each cell. Since the GO term activity score is account for the expression of multiple genes, it is expected to be more reliable than the scRNA-seq dataset, and therefore, benefit the downstream analysis, like cell clustering, etc. Again, CMC model is applied here to jointly consider the impacts of both cell and gene factor classes on scRNA-seq data.

**Fig. 6.**
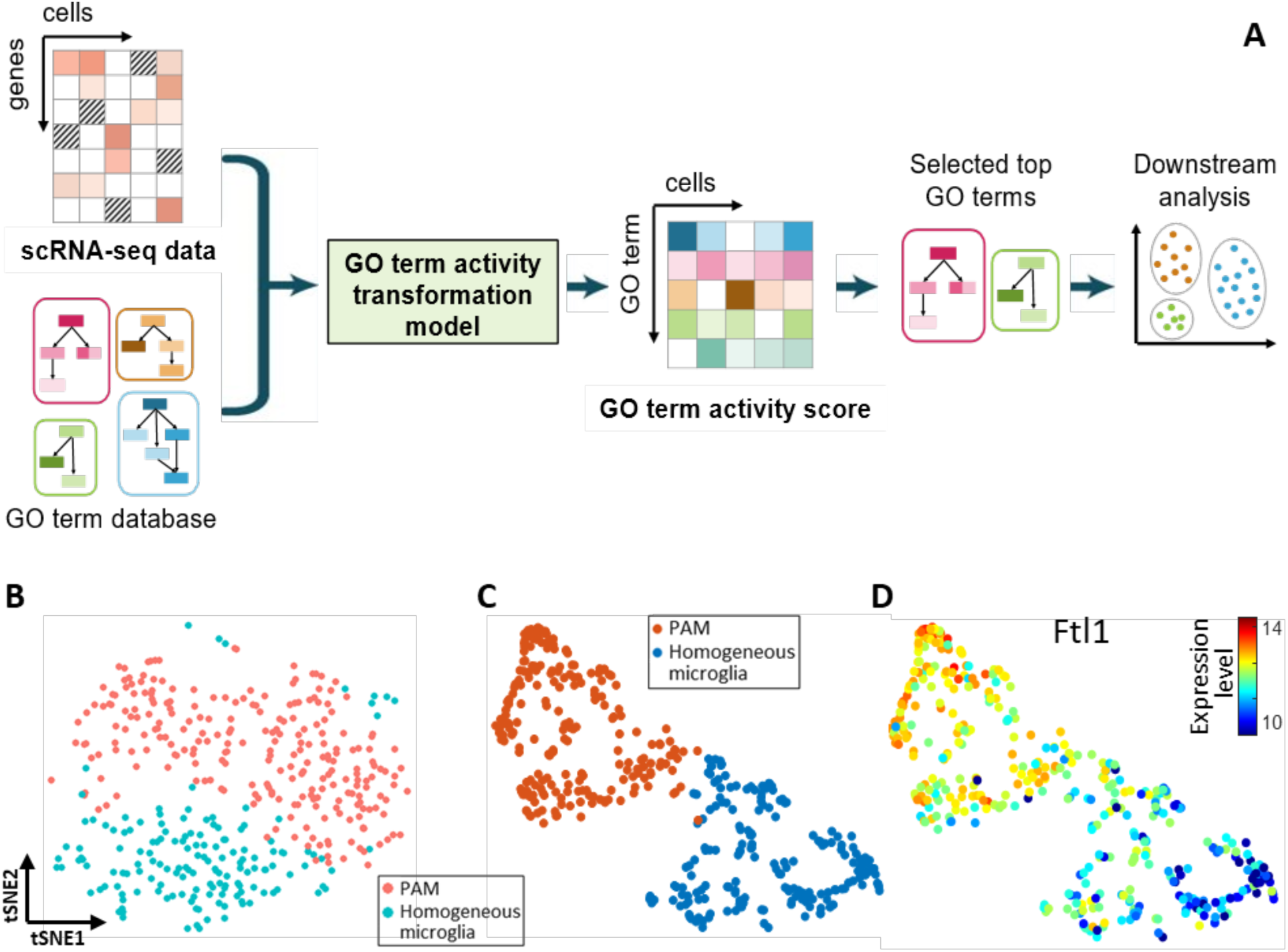
(A) Introduction of the GO term activity transformation. By incorporating pri-knowledge of GO terms, the scRNA-seq data is transformed into GO term activity scores. In the transformed data, each row represents a GO term and each column corresponds to a cell. The activity score indicates the level of activity of a specific GO term within a particular cell. The transformed data can be used for downstream analysis, such as clustering, etc. (B-D) Application to a real dataset (*24*). (B) Clustering analysis results based on scRNA-seq data, which barely separated PAM from homogenous microglia. (C) Clustering analysis results based on GO term activity score. (D) The expression pattern of PAM’s marker, Ftl1.

We expected our GO term activity score is more powerful in identifying cell heterogeneity, because 1) the GO term activity score data is more robust as each value takes into account the expressions of multiple gene; 2) the gene-level expression bias can be modeled and captured by CMC model, therefore, genes expressed homogeneously in the cell population will have less contribution to the final activity scores.

To validate the effectiveness of our model, we apply it to the dataset of (*24*) (Fig. 6B). Reference (*24*) confirmed that PAM is a novel type of microglia that are different from homogenous microglia. However, as shown in Fig. 6B, the clustering analysis based on gene-level expression data barely separated PAM from homogenous microglia.

Here we transform the scRNA-seq data into the GO term activity score, and redo the clustering analysis. As shown in Fig. 6C, PAM and homogenous microglia are well separated. Note that there is a narrow bridge between these two groups. These cells are likely cells in the transition state. Fewer transition cells are shown here. This is probably because the scRNA-seq technology is basically taking a snapshot of the cells at certain time point. Compared with the cells that are in a stable state (e.g., PAM and homogenous microglia), the cell in a rapid transition state is less likely to be captured, and consequently, less number of transition-state cells. Our model also provide the opportunity to directly inspect the activity level of any GO term in each cell. Fig. 6D shows the expression level of a marker of PAM cells, which demonstrate a progressive pattern of change from PAM to homogenous microglia.

The CMC model is designed to jointly consider the impacts of all factors among data. Its solution is simple but elegant and can be solved efficiently. It’s flexible to handle data associated with multiple factors, with binary/integer values, and values attached by real-valued weight.

CMC also allows data with missing values. The CMC model has demonstrated its effectiveness in five applications, and especially, has led to genuine biological discoveries in identifying the TF that function in oligodendrocyte lineage development. Consider that the real bioglocal data are often affected by numerous hidden factors, we believe the CMC model will show its broad value in science and engineering.

## Notes

### Competing Interest Statement

The authors have declared no competing interest.

