## Supplementary for "An Efficient and Principled Model to Jointly Learn the Agnostic and Multifactorial Effect in Large-Scale Biological Data"

**This PDF file includes:**

Materials and Methods
Figs. S1 to S14
Table S1

### Materials and Methods

#### 1 Application to scRNA-seq data normalization

1.1 Introduction

scRNA-seq is a technology to profile the RNA molecule quantity in each cell. It provides us with a powerful tool for revealing exciting biological insights in single-cell resolution (24,25). In general, the scRNA-seq isolates the biology sample into single cells. Then for every single cell, the mRNA molecules in its lysis are captured (by poly[T] priming) and are reverse transcribed into cDNA. Subsequently, the tiny amount of cDNAs are amplified by techniques like PCR. Finally, the cDNA fragments, resulting from the fragmentation, are sequenced (24). After aligning the sequenced fragments to the referent genome, raw count, which records the number of fragments of each gene in every single cell, is obtained.

Current scRNA-seq technologies can be roughly divided into UMI-based technologies and full-length technologies. Full-length technologies convert the entire mRNA into cDNA and sequence them after fragmentation and amplification. The UMI-based technologies, on the other hand, only sequence part of the mRNA. They sacrificed the full-length coverage to increase the throughput of scRNA-seq library generation by one to three orders of magnitude and dramatically reduce the sequencing cost (26,27). These methods allow us to tag a unique molecular identifier (UMIs) to each mRNA molecule. Therefore, each mRNA will be correctly identified and counted even though it has many duplicates after the amplification step (28). The UMI-based methods count the absolute numbers of mRNA molecules and have advantages in cost (29). Full-length technologies can cover the whole genome sequence. And compared with UMI-based technologies, they are more sensitive, especially for those lowly expressed genes (19).

The raw counts of scRNA-seq are susceptive to factors like cell's library size, gene's GC content, and gene length, etc. The cell's library size directly impacts the total read counts of a cell. The GC content of a gene's sequence will impact the efficiency of PCR amplification. As for gene length, compared with short genes, the longer ones will result in more cDNA fragments and are more likely to be sequenced during the RNA-sequencing. The first one is the technical factor impacting every single cell, and the last two are inherent to the gene. We categorize the first one as cell factor and the last two as gene factor respectively. Besides these two factors, we found that, in the full-length sequencing technologies (such as Smart-seq2), the "cDNA length" is also an important factor that has a non-negligible impact on the raw counts.

We developed a framework to normalize the raw count of full-length-based scRNA-seq technologies. In the framework, the CMC is applied to jointly infer the factors mentioned above (cell, gene, and cDNA-length factors) and then to normalize out the cell and cDNA-length factors. The heterogeneity among genes is kept in the normalized data because it includes information of gene's average expression levels across all cells, which might be of interest to the users. Users can choose to normalize out the gene factor as well if they want.

1.2 The cDNA-length factor

The cDNA length refers to the sequence length of a cDNA molecule after its reverse transcription from the template mRNA during the scRNA-seq library preparation. Compared with a short cDNA molecule, a longer one tends to generate more cDNA fragments after the fragmentation procedure and is expected to have more fragments to be successfully sequenced in the final step of sequencing, resulting in a higher read count.

Typically, the length of a cDNA molecule is equal to that of the mRNA that it reverse-transcribes from, if an entire mRNA molecule is successfully reverse-transcribed to cDNA (the lengths of primers are not considered). However, incomplete reverse transcription may occur and result in truncated cDNAs, which are shorter than they should be (*17*). As a consequence, fewer fragments are expected to be generated and successfully sequenced. Fig. S 1 illustrates normal and incomplete reverse transcription (take the protocol of Smart-seq2 as an example). In the normal case, the oligo(dT) primer attaches to the mRNA's poly-A tail, where the reverse transcription starts from. After fully reverse-transcribing the mRNA template, extra nucleotides, Cs, are added, and the ISPCR primer will be attached to the 5' cap. In this case, the mRNA will be fully reverse-transcribed (Fig. S 1A). However, in some abnormal circumstances, the oligo(dT) primer may wrongly attach to the mRNA's coding sequence, and reverse transcription will start from that position. Besides, extra Cs and ISPCR primer may also improperly bind to the mRNA's coding sequence, resulting in an early stop of reverse transcription. As a result, only a part of the mRNA sequence is successfully reverse transcribed, as shown in Fig. S 1B.

The phenomenon of incomplete reverse transcription is also revealed by the sequenced fragments, which is the outputs of a sequencing machine. By aligning a sequenced fragment to the reference genome, we precisely identify the part of the mRNA sequence that the fragment was initially reverse-transcribed from. We found that, in some cases, the fragments of an individual gene in one cell were all originally from a small part of the gene's mRNA sequence. Fig. S 2A shows the alignment results of fragments corresponding to ERCC-00116 in a cell. It can be seen that all these fragments are squeezing in a small region of ERCC-00116's sequence, implying all these sequenced fragments initially come from a small part of an mRNA molecule. Such observation can be explained by the incomplete reverse transcription, where only this small part of mRNA is successfully "copied" for the downstream amplification and sequencing. By design, during the library preparation, an entire mRNA molecule is expected to be reverse-transcribed to cDNA and to be sequenced after amplification and fragmentation. As a result, the sequenced cDNA fragments are expected to cover the whole mRNA. Fig. S 2B shows a typical case where the sequence reads align to the entire region of the gene's reference sequence. These phenomena can also be observed in other genes (Fig. S 2C and Fig. S 2D).

A concern may arise with the causality between the number of fragments and the reference sequence's region covered by the fragments. Specifically, it may be the small number of sequenced fragments that results in the narrow coverage of the reference region by the fragments. A piece of direct evidence to address this concern is the ISPCR primers. ISPCR primers are used and only used to attach the ends of the reverse-transcribed cDNA (16) (Fig. S 1). For the cases shown in Fig. S 2A and Fig. S 2C, we indeed found the fragments whose 5' or 3' end aligning to the end of the narrowly covered region is attached by ISPCR primer, implying that the fragments already cover both the 5' and 3' ends of the cDNA they originated from. Besides, statistical tests are also performed for cases shown in Fig. S 2A and Fig. S 2C, to test whether the narrowly covered regions are due to the small number of fragments. For this purpose, we randomly permuted the location of each fragment for $10^9$ times, and computed the lengths of the covered regions. The null distribution is built by fitting the distribution of covered region lengths obtained from the $10^9$ permutations. We then accessed the p-value of the observed covered region length. The p-values for Fig. S 2A and Fig. S 2C are both smaller than $10^{-300}$, rejecting the null hypotheses that the small length is due to the small number of sequenced fragments.

Though the sequence length of each reverse-transcribed cDNA molecule was not recorded during the sequencing, it could be roughly inferred by aligning the corresponding fragments to the referent sequences and calculating the length between the first and the last nucleotide covered by the fragments. However, there is neither enough information for us to tell the original cDNA molecule for each sequenced fragment, nor the set of fragments that originate from the same cDNA molecule. Therefore, for the fragments corresponding to a gene (which could be told by the alignment results), we do not distinguish which cDNA molecule they are coming from. Instead, we infer the longest possible length of these fragments' original cDNA molecule. As a result, a single cDNA length value is inferred for all fragments of a gene in a cell.

The product of incomplete reverse transcription is truncated cDNA, which is shorter than the normal one and will generate fewer fragments in the following steps. As a result, the corresponding mRNA tends to have smaller read counts. We observed such a tendency in scRNA-seq datasets from multiple sources (18, 19).

Fig. S **3** shows the tendency of a gene's read counts across cells, as a function of its corresponding cDNA length. Noted that "ERCC-00116" is a spike-in RNA, which is manually added to each cell lysis evenly as a control probe. Its read counts are expected to be similar across all cells. This is true when the mRNA is completely reverse transcribed (in such cases, the corresponding cDNA lengths are equal to the RNA length, which is 1991). Fig. S 3A implies that the read counts tend to be smaller as the reverse-transcribed cDNA becomes shorter. We also observed this phenomenon for other genes (Fig. S 3B and Fig. S 3C).

cDNA length factor was regarded as an additional factor during the normalization procedure. It was not categorized into the gene factor as the gene length did, because the corresponding cDNA length of the same gene varies across cells.

Besides, preliminary evidence shows that the cDNA length factor is not independent of other facts, such as the cell factor (Fig. S 4), highlighting the necessity of jointly considering the gene, cell, and cDNA length factors.

In summary, incomplete reverse transcription may occur during the RNA sequencing, resulting in shorter cDNAs and, consequently, smaller read counts than they should be. It is necessary to take the cDNA length into account when normalizing the scRNA-seq raw counts. What's more, as the cDNA length factor is not independent of other factors, the cDNA length factor, along with other factors, should be considered jointly. As the CMC model consider multiple factors jointly, we shall apply it to the scRNA-seq data normalization task.

#### 1.3 Framework

The cDNA length was divided into 15 levels according to the interval it lied in: [1, 200], [201, 300], [301, 400], [401, 500], [501, 600], [601, 700], [701, 1000], [1001, 1500], [1501, 2000],

[2001, 2500], [2501, 3000], [3001, 3500], [3501, 4000], [4001, 4500], and [4501,+∞). And the scRNA-seq read counts data are stored in a three-dimensional (3D) contingency tensor, with each dimension corresponding to the gene, cell, and cDNA length, respectively. A value in the tensor represents the raw count of each gene in each cell with its cDNA length lying in a specific cDNA length interval. Given the 3D tensor, the CMC model was then applied to estimate the expected value of each entry with joint consideration of the gene, cell, and cDNA length factors. After the expected values were estimated, the raw count and the expected values in the 3D tensors were summed back to two-dimensional (2D) tensors respectively along the cDNA length dimension. Finally, the read counts were normalized via dividing the raw count by its expectation and taking a logarithm transformation.

As mentioned in the main text, we regarded the 0s as missing values when inferring the gene/cell/cDNA-length factors. The 0 values in the raw count of scRNA-seq are probably due to dropout events during the RNA sequencing. Such 0s do not necessarily imply that the corresponding genes have no expression. For example, dropout events happen when the gene's mRNAs fail to be captured during the reverse transcription procedure. We ignored the 0s while modeling the heterogeneities among genes, cells, and cDNA lengths, and regard them as missing values. The CMC model is compatible with missing values and was applied to infer the susceptivities values for each gene, cell, and cDNA-length category, and the expectation values of each entry conditional on the inferred susceptivities of cell factor and cDNA-length factor.

#### 1.4 More experimental results

One of the criteria we used to evaluate the normalization performance is the variance of each gene's expression across a set of homogeneous cells. That is, to measure how much of the variance due to noise (both sequencing noise and biological noise) is removed after the normalization. In addition to the experimental results shown in the main text, we provided results on two more datasets that are from another source (*19*) and of different cell types.

The two datasets are the scRNA-seq data of 4,381 homogeneous microglia cells and 1,338 homogeneous oligodendrocyte cells. Fig. S 5 (the first row) shows each gene's variance change after normalization. Normalization using the CMC model reduces the variance of most genes to about half of their original variance. For comparison, Fig. S 5 (the second row) shows the variance changes after being normalized by a popular normalization method, total count (TC) (*5*), which considers the cell factor separately from the gene factor. It can be seen that, for the TC method, the variances reduce dramatically for genes with very small zero rates (corresponding to a small part of extremely highly expressed genes). However, except for this small part of genes, the rest of the genes' variances are only slightly changed. This is because the total count of each cell is dominated by a small number of extremely highly expressed genes. Normalizing out the cell factor without jointly considering such heterogeneity among genes will result in the normalization result favoring those highly expressed genes. Fig. S 5 (the third row) compares the variance of each gene after normalizing by CMC model and by TC method respectively. It confirmed that the TC method emphasized more on a small part of genes with extremely large values. Fig. S 5 (the last row) summarizes the genes' variance change after the normalization performed by the CMC model and other normalization methods: TC, TMM(*7*), RUV(*8*), UQ(*6*), SCnorm(*9*), Scran(*10*), and Census (*11*). It can be seen that the CMC model outperforms all the other methods in terms of reducing genes' variances.

### 2 Application to driving TF identification

Driving TF identification takes as input a set of co-expressed genes and identifies the driving TFs that are responsible for the co-expression of this set of genes. The framework of this application is shown in Fig. 4 and introduced in the main text. In short, the TF-gene binding state data is firstly derived from ChIP-seq databases and stored in a 2D tensor. Then the CMC model is applied to infer the binding affinities of each TF/gene, as well as the binding probability of each TF-gene pair. Finally, the driving TFs are identified by comparing each TF's observed binding state with the input co-expressed gene set and the corresponding binding probability.

In this section, we will describe the details of the implementation of this framework, which includes the following key modules: TF-gene binding state database, cell-type-specific TF-gene binding probability, and the statistical test.

#### 2.2 Module 1: TF-gene binding state database

The TF-gene binding state tensor indicates whether a TF is binding to a gene. The tensor is two-dimensional, in which one column represents a gene, and one row corresponds to a TF attached by cell-type property. Value 1 in an entry $(i, j)$ indicates that the $TF_i$ binds to and consequently regulates the $gene_j$ in the corresponding cell type. Value 0 indicates $TF_i$ does not bind to $gene_j$ in that cell type. The 2D tensor collects the binding states between different TFs and genes in various cell types. It serves as a knowledge base in the driving TF identification model. It's the binding state matrix that the model relies on to identify the TFs that significantly "favor" the co-expressed gene list. A binding matrix of good quality is critical for the identification task.

Datasets from ChIP-seq (chromatin immunoprecipitation followed by sequencing) experiments were selected to extract the TF-gene binding state. Each dataset records the result of one ChIP-seq experiment. ChIP-seq experiments can be performed to detect the genome-wide binding sites of an individual TF in a specific tissue/cell type. It is wildly used to detect the TF-DNA interactions in vivo. ChIP-seq datasets were selected for the following considerations: 1) ChIP-seq data of a TF can accurately characterize the TF's genome-wide binding sites (4). 2) ChIP-seq data can capture the cell type-specific binding states between TFs and genes. 3) There are thousands of ChIP-seq datasets for different TFs in different cell types (30, 36-39).

We extracted the TF-gene binding states from UniBind's TF-DNA interactions Database (30), which comes from 9654 ChIP-seq experiments across nine species, for its high standard in terms of data quality. UniBind uses two quality control metrics to ensure the quality of inputted ChIP-seq datasets. 1) It expects the high-quality ChIP-seq peak datasets to be enriched for the TF binding motif known to be bound by the ChIP'ed TF, and filters out the ChIP-seq datasets not satisfying this metric. 2) It filters out the datasets where the predicted TFBSs did not show a significant enrichment around the summits, as it expects the high-quality ChIP-seq peaks to be enriched for TFBSs close to their summits. In addition, UniBind combines both computational (via TF motif information) and experimental (via ChIP-seq datasets) evidence to find the high-confidence TF-gene interactions.

A TF regulates its targeted gene's expression by binding to either enhancer or promoter of the targeted genes. However, converting the TF-DNA interactions to the TF-gene binding states is not trivial for the following reasons: 1) although promoters are typically immediately adjacent to the gene's transcriptional start site (TSS) in DNA sequence, the exact length of each gene's promoter can often only be defined experimentally; 2) The enhancer can be located upstream or downstream up to 1 Mbp away from the TSS of its targeted gene, and there is no accurate map linking enhancers to their target genes (40); 3) not all TF's bindings are functional (41). A score is therefore needed to quantify how likely a gene is regulated by a TF. While estimating the score, the effect a TF binding site has on the expression of its targeted gene is assumed to decay exponentially with the genomic distance between the binding site and the TSS (42). While TFs can also regulate genes by binding to genes' enhancers, it's almost impossible to incorporate such regulation into the database, as discussed above (43). Therefore, we only consider the TF binding sites located around the TSS.

To calculate the binding score between any TF-gene pair, a distance weight $w$ representing the regulatory influence of a locus at position $k$ on the TSS of gene $j$, is firstly defined (4):

$$w_{kj} = \frac{2e^{-\mu d_{kj}}}{1 + e^{-\mu d_{kj}}}$$

where $d_{kj} = |k - t_j|$ is the distance between nucleotide position $k$ and the position of gene $j$'s TSS at $t_j$. $\mu = \frac{ln3}{\Delta}$ is the parameter to determine the decay rate of the weight; $\Delta$ is a predefined decay distance, corresponding to the distance that $w_{ik}$ decay to 0.5. The binding score between TF $i$ and gene $j$ is:

$$S_{ij} = \max_k w_{kj} s_{ki}$$

where $s_{ki} \in \{0,1\}$ is the signal of whether TF $i$'s binding at genome position $k$. $S_{ij} = [0,1]$.

We found that, for ChIP-seq datasets from different labs, their TF-DNA interaction counts vary dramatically, even for the same TF in the same cell types. Such difference is probably due to the batch effects. We assigned different decay distances $\Delta$ to roughly control the impact of the batch effect. The $\Delta$ was set to 250, 500, and 1000 for datasets with TF-DNA interaction counts less than 256, between 256 and 1024, larger than 1024, respectively.

The final TF-gene binding state tensor is illustrated in Fig. S 6, in which one row corresponds to a ChIP-seq dataset, and one column represents a gene. In the tensor, TFs with an identical name but from different datasets are regarded as different TFs and are represented by different rows. This is because the datasets of TFs with an identical name are the results of ChIP-seq experiments that may be conducted on various cell types, and the TF-gene binding states are cell type-specific. TFs with an identical name will be jointly considered in the final step of driving TF identification. For each dataset, its corresponding TF and cell type are attached. This tensor has 3072 rows (datasets) and 24528 columns (genes).

### 2.3 Module 2: Cell type-specific TF-gene binding probability

The binding probability matrix is inferred from the TF-gene binding state matrix. Value in entry $(i, j)$ is the probability that the $j^{th}$ gene is bound by the TF of the $i^{th}$ ChIP-seq dataset in the corresponding cell type, conditional on the TF's/gene's binding affinities in the cell type. The probability matrix is used to compare with the binding state matrix so as to identify the TFs that are significantly enriched among the inputted gene set.

The binding probability is often cell type-specific, due to the cell type-specific regulation (31-35). Evidence from the TF-gene binding state tensor also supports this statement (Fig. S 7). Fig.

S 7 shows that the binding affinities of a gene are close in similar cell types/tissues, but could be quite different in distinct cell types/tissues. Therefore, a cell type-specific TF-gene binding probability is needed.

To learn the cell type-specific TF-gene binding probability from the binding state tensor, a straightforward method is regarding the cell type as a third factor and storing the binding state information in a three-dimension (TF, gene, and cell type) tensor, then applying the CMC model to infer the cell-type-specific binding probability. However, the ChIP-seq datasets that have been generated are far from covering all TFs and all cell types (35). For many cell types/tissues, only a limited number of ChIP-seq datasets are available. For these cell types/tissues, it's hard to confidently estimate the TF-gene binding probability given the small number of datasets.

We developed a CMC-based framework to confidently estimate the cell type-specific TF-gene binding probability (Fig. S 8). Basically, for each cell type, the ChIP-seq datasets corresponding to this cell type were used as seeds to identify the ChIP-seq datasets coming from similar cell types. The seeds and the identified ChIP-seq datasets would jointly contribute to the inference of the TF-gene binding probability for this specific cell type. For convenience, we call the ChIP-seq datasets coming from similar cell types of the seeds as seeds' "related ChIP-seq datasets". To identify the related ChIP-seq datasets for each cell type, we first identified from the seeds a set of representative genes, which represents the binding characteristic of this cell type. Given the representative genes, the related ChIP-seq datasets were then identified.

The representative genes of a cell type are the genes that are more frequently bound by various TFs in this cell type than in other cell types (the background cell types). These genes contribute to the binding characteristics of this cell type. As TF regulates the expression of its target genes by binding to them, these genes are likely to be DEGs in this cell type compared with the background cell types. To identify the representative genes, the CMC model was reused again. The identification module is similar to that of driving TF identification (Figure 4A), except that the input is the IDs of the seeds (i.e., the ChIP-seq datasets of a specific cell type). In such cases, given the binding genes of all ChIP-seq experiments performed on various cell types, the module identifies the genes that are enriched in the ChIP-seq of a specific cell type. To this end, for each gene, its observed binding states among the seeds were compared with the corresponding binding probabilities, and a p-value was assessed. To infer the binding probability, the CMC model was applied due to the existence of heterogeneities across genes and datasets. Unlike the driving TF identification, we do not need to consider the cell-type-specific binding probability, because the binding heterogeneity between cell types is the information that we rely on for the identification. If we regard the "cell types" as an additional dimension then the CMC model would model and remove the impact of the heterogeneity of cell types, and the desired cell type-related datasets would not be identified.

Given the representative genes of a specific cell type, we then detected the related ChIP-seq datasets, in which the representative genes are frequently bound. The assumption behind this is that the seeds and their related ChIP-seq datasets have similar binding characteristics. Specifically, their representative genes are highly overlapped. The module of related ChIP-seq datasets identification is also similar to Figure 4A, except that the input is the representative genes. In such a case, the module identified the ChIP-seq datasets that the representative genes are bound more frequently than expected.

Given the identified related ChIP-seq datasets, we then combined them with the seed datasets to jointly infer the cell type-related binding probability matrix. Specifically, for each cell type, its corresponding seeds and identified related ChIP-seq datasets were used to infer the cell-type-specific gene factor, during which, the binding scores of each dataset are multiplied by a weight reflecting how well the dataset represents the binding characteristics of the cell type. The weights are transformed from the p-values of the related datasets identified. Fig. S 11 showed the correlation coefficient of the estimated genes' binding affinities between any two cell types. It can be seen that similar cell types have relatively higher correlation coefficients compared with those among distinct cell types. The results are consistent with our expectation that similar cell types have similar genes' binding affinities.

Given the genes' cell-type-specific binding affinities, we then inferred the cell-type-specific binding probabilities for the "TF-gene binding state tensor" shown in Figure 4A. For each cell type, the CMC model was applied to the whole tensor with genes' cell-type-specific binding affinities fixed, and the cell-type-specific binding probabilities are estimated for this cell type.

### 2.4 Module 3: Statistical test

To detect the driving TFs of the inputted co-expressed gene set, for each TF, we checked whether it binds to the co-expressed gene set more frequently than by random. Specifically, let $x_1, \ldots, x_n$ be the binding states of the examined TF and the $n$ co-expressed genes, and $P_1, \ldots, P_n$ are the corresponding binding probabilities conditional on the TF's and genes' binding affinities; we assess the p-value of the score:

$$S_{obv} = \sum_{i=1}^{n} \omega_i x_i$$

under null hypothesis

$$S = \sum_{i=1}^{n} \omega_i b_i$$

where $b_i$ is the Bernoulli variable with success probability $P_i$; $\omega_i$ is a predefined weight attached to $b_i$. We set $\omega_i = \frac{1}{\sqrt{P_i(1-P_i)}}$ to guarantee that each weighted Bernoulli variable has the same variance and equally contributes to $S$.

As we carefully considered the binding affinity of each gene, $\{P_i\}$ are typically different, making it extremely hard to compute the exact p-values. Therefore, as mentioned in the main text, we utilized saddle point approximation to provide an accurate and efficient alternative, i.e., $p(S_{obv}) = 1 - \hat{F}(S_{obv})$, where $\hat{F}(S_{obv})$ is the approximated CDF. It is given by (44):

$$\hat{F}(S_{obv}) = \begin{cases} \Phi(\hat{\alpha}) + \phi(\hat{\alpha})\left(\frac{1}{\hat{\alpha}} - \frac{1}{\hat{\beta}}\right) & \text{for } x = \text{Exp}(S) \\ \frac{1}{2} + \frac{K'''(0)}{6\sqrt{2\pi}\, K''(0)^{\frac{3}{2}}} & \text{for } x \neq \text{Exp}(S) \end{cases}$$

where $K(S) = \log(M(S))$ is the cumulant generating function; $M(S)$ is the moment generating function of the distribution of $S$; $\hat{\alpha} = \text{sgn}(\hat{y})\sqrt{2(\hat{y}S_{obv} - K(\hat{y}))}$ , $\hat{\beta} = \hat{y}\sqrt{K''(\hat{y})}$, and $\hat{y}$ is the solution to $K(\hat{y}) = S_{obv}$.

After the statistical test, for TFs with identical names, only the most significant one was reported.

### 3 Application to cancer-associated gene identification

#### 3.1 Introduction

Major international projects like The Cancer Genome Atlas (TCGA)(45) and the International Cancer Genome Consortium (46) created comprehensive mutation data across all major cancer types. Such data include the identified genomic mutations that occur in cancer patients obtained via the whole-genome sequencing for their matched tumor-normal cells. Mathematical analysis of this mass data to identify the mutated genes that drive cancer pathogenesis is of huge help to understand its mechanism. The basic idea is to identify the genes with significantly more mutations than expected among the patients suffering a specific type of cancer.

However, estimating the expected mutation rate is not trivial due to the mutational heterogeneities in cancer (12). Specifically:

1) The mutation rates vary markedly across and within cancer types. Took the data from TCGA for example, the median mutation frequency varies by more than 1,000-fold across cancer types. And within a cancer type, the patients' mutation frequencies are also quite different. For example, the mutation rate in the lung adenocarcinoma cancer ranged across 0.1–100/Mb (Mb: million base pairs);

2) The mutation rate shows diversity among different mutation categories (CpG transversions, C:G transitions, A:T transversions, etc). For example, the "CpG transversions" have a much lower mutation rate than other categories (Fig. S 12).

3) The mutation rates markedly vary across the genome regions. The diversity of gene characteristics (especially the expression levels and replication time during the cell cycle) may be responsible for such genome regional heterogeneity, and the gene characteristics are cell-type-specific (12, 47).

We summarized the mutational heterogeneities mentioned above into three aspects: heterogeneities of patient factor, mutation category factor, and gene factor respectively.

The heterogeneities of these three factors are not independent of each other. As it was shown in Fig. 1B in the main text, the mutation rates of any two factors are twisting together.

Ignoring the heterogeneities of one or more factors might lead to awkward results. For example, MutSig1.0 ignores the heterogeneity across genes. As a result, unreasonable results often occur (12). For example, with larger sample sizes, the number of falsely significant genes grows rapidly, which is opposite to the expectation. Larger sample sizes are expected to have more power both to detect the true driving genes (sensitivity) and distinguish them from the false-positive genes (specificity). More specifically, MutSig1.0 identified a total of 450 significant genes (false-discovery rate $q < 0.1$), at least 257 of them are highly likely to be false discoveries according to their biological function or genomic properties. MutSigCV (12), on the other hand, took mutational heterogeneities in all three factors into account. It does reduce the false positive in some cases. For example, when MutSigCV is applied to the squamous cell lung cancer example, it only identified 11 significant genes. However, as it addressed the factors independently and estimated the joint mutation rate by the product of marginal relative rates of the three factors, MutSigCV identifies many false-positive genes in some cancer types. For example, it identifies about 1498 significant genes for lung adenocarcinoma cancer and more than 450 genes for the other three cancer types. These two example methods highlight the necessity of jointly considering the heterogeneities of all three factors.

#### 3.2 Framework

We developed a framework to identify the cancer-associated genes of a specific cancer type, where the CMC model is applied to jointly model the heterogeneities among patients, genes, and mutation categories. As shown in Fig. S 13, the mutation counts are stored in a three dimensions tensor, where the three dimensions correspond to gene factor, patient factor, and mutation category factor, respectively. The susceptivity of each patient, gene, and mutation category was then jointly inferred, and the expected number of mutations conditional on the susceptivities of the corresponding patient, gene, and mutation category are estimated. The framework takes as input the IDs of patients that suffer from a specific type of cancer and identified the genes associated with this type of cancer. To this end, for each gene, a statistical test is performed by comparing its mutation counts among the patient suffering this type of cancer and the corresponding expectations.

#### 3.3 Implementation details

The mutation datasets of 9298 samples across 31 cancer types were downloaded from (30). In the raw data, each item records a single mutation and contains corresponding information such as cancer type, patient ID, gene, referent sequence, mutated sequence, and mutation types. Nonsilent mutations were selected from the data and grouped into seven categories: CpG transitions, CpG transversions, C:G transitions, C:G transversions, A:T transitions, A:T transversions, and null and indel mutations. The mutation data were stored in a three-dimensional tensor, with each dimension corresponding to patient, gene, and mutation category, respectively. The tensor was then binarized. During the quality-control procedure, patients with no more than 10 mutations (across all genes and mutation categories) and genes with no more than 10 mutations (across all patients and mutation categories) are filtered out. As a result, a tensor with 8544 patients, 16277 genes, and 7 mutation categories was obtained.

Given the 3D tensor, the CMC model was applied to infer the mutation susceptivities of each patient, gene, and mutation category, and the mutation probability of each entry conditional on the inferred mutation susceptivity of its corresponding patient, gene, and mutation category.

Then, for the inputted patient IDs that suffered a specific type of cancer, the framework identifies the associated genes by checking whether a gene has significantly more mutations than expected among these patients. To this end, saddle point approximation was again applied to access the p-value for each gene, the same as it was in the application of driving TF identification.

### 4 The CMC model

4.1 The basic CMC model for three-dimensional binary tensor

We first introduce the basic model for a three-dimensional $(N \times M \times Q)$ binary contingency tensor. The task is to infer the probability distribution of each entry given the marginal totals in each dimension.

We assume that each entry of the contingency tensor is independent, and follows the Bernoulli distribution with probability $P_{ijk}$ for all $i \in \{1,...,N\}$, $j \in \{1,...,M\}$, and $k \in \{1,...,Q\}$. $P_{ijk}$ is inferred by maximizing the entropy under the marginal expectation constraints:

$$\max_{\vec{P}} \; -\sum_{i=1}^{N}\sum_{j=1}^{M}\sum_{k=1}^{Q}\left[P_{ijk}\ln P_{ijk} + \left(1-P_{ijk}\right)\ln\left(1-P_{ijk}\right)\right] \qquad \text{(S4.1)}$$

$$s.t. \quad \sum_{k=1}^{Q}\sum_{j=1}^{M} P_{ijk} = n_i, \text{for } i = 1, \dots, N$$

$$\sum_{k=1}^{Q}\sum_{i=1}^{N} P_{ijk} = m_j, \text{for } j = 1, \dots, M$$

$$\sum_{j=1}^{M}\sum_{i=1}^{N} P_{ijk} = q_k, \text{for } k = 1, \dots, Q$$

where $n_i$, $m_j$, and $q_k$ are the marginal totals of the $i^{\text{th}}$, $j^{\text{th}}$, and $k^{\text{th}}$ individual in the three dimensions, respectively.

To solve Eq. (S1), Lagrange multipliers $\{r_i\}$, $\{w_j\}$, and $\{u_k\}$ are introduced:

$$\max_{\vec{P},r,w,u} -\sum_{i=1}^{N}\sum_{j=1}^{M}\sum_{k=1}^{Q}\left[P_{ijk}\ln P_{ijk} + \left(1-P_{ijk}\right)\ln\left(1-P_{ijk}\right)\right] + \sum_{i=1}^{N} r_i\left(\sum_{k=1}^{Q}\sum_{j=1}^{M} P_{ijk} - n_i\right) + \sum_{j=1}^{M} w_j\left(\sum_{k=1}^{Q}\sum_{i=1}^{N} P_{ijk} - m_j\right) + \sum_{k=1}^{Q} u_k\left(\sum_{j=1}^{M}\sum_{i=1}^{N} P_{ijk} - q_k\right) \qquad \text{(S4.2)}$$

Taking the derivative with respect to $\{P_{ijk}\}$ and setting it to 0:

$$-\left[\ln P_{ijk} + 1 - \ln\left(1-P_{ijk}\right) - 1\right] + r_i + w_j + u_k = 0 \qquad \text{(S4.3)}$$

The optimal probability form for each entry is obtained by reorganizing Eq. (S4.3):

$$P_{ijk} = \frac{e^{r_i+w_j+u_k}}{e^{r_i+w_j+u_k}+1} \qquad \text{(S4.4)}$$

Having obtained the probability distribution from Eq. (S4.4), the susceptivities, i.e., the parameters $\{r_i\}$, $\{w_j\}$ and $\{u_k\}$, can be learned through maximum likelihood estimation:

$$\max_{r,w,u} \sum_{i=1}^{N}\sum_{j=1}^{M}\sum_{k=1}^{Q} \ln\left[x_{ijk}\left(\frac{e^{r_i+w_j+u_k}}{e^{r_i+w_j+u_k}+1}\right) + (1-x_{ijk})\left(\frac{1}{e^{r_i+w_j+u_k}+1}\right)\right] \qquad \text{(S4.5)}$$

where $x_{ijk} \in \{0,1\}$ is the observed value in the entry $(i, j, k)$ of the tensor. Taking the derivative with respect to $\{r_i\}$, $\{w_j\}$, and $\{q_k\}$, and setting them all to 0, we have:

$$\begin{cases} \sum_{j=1}^{M}\sum_{k=1}^{Q}\frac{e^{r_i+w_j+u_k}}{e^{r_i+w_j+u_k}+1} = n_i & i = 1,\dots,N \quad (S4.6a) \\ \sum_{i=1}^{N}\sum_{k=1}^{Q}\frac{e^{r_i+w_j+u_k}}{e^{r_i+w_j+u_k}+1} = m_j & j = 1,\dots,M \quad (S4.6b) \\ \sum_{i=1}^{N}\sum_{j=1}^{M}\frac{e^{r_i+w_j+u_k}}{e^{r_i+w_j+u_k}+1} = q_k & k = 1,\dots,Q \quad (S4.6c) \end{cases} \qquad \text{(S4.6)}$$

where $n_i = \sum_{j=1}^{M}\sum_{k=1}^{Q} x_{ijk}$ , $m_j = \sum_{i=1}^{N}\sum_{k=1}^{Q} x_{ijk}$ , and $q_i = \sum_{i=1}^{N}\sum_{j=1}^{M} x_{ijk}$ are the marginal totals of the tensor.

There is infinite number of solutions for Eq. (S6), even the number of variables ($N+M+Q$) is equal to that of equations. This is because $\sum_{i=1}^{N} n_i = \sum_{j=1}^{M} m_j = \sum_{k=1}^{Q} q_k$, i.e., at least two equations in the Eq. (S6) are the combinations of the others. However, the value of $r_i + w_j + u_k$ for all $i \in \{1,...,N\}$, $j \in \{1,...,M\}$, and $k \in \{1,...,Q\}$ are always fixed. And consequently, $P_{ijk}$, the function of $r_i + w_j + u_k$, has a unique solution (Theorem 4.1). Besides, the ratio between any two variables of the same factors is also fixed (Theorem 4.2).

We proposed an iterative strategy (Algorithm 1) to find a solution to Eq. (S4.6), in which, variable sets $\{r_i\}$, $\{w_j\}$ and $\{u_k\}$ are updated sequentially. And during the update of each variable set $\{r_i\}$, $\{w_j\}$ or $\{u_k\}$, variable elements within each set can be updated parallelly, given the fact that the variable elements within each set are independent if the other two variable sets are fixed. For each single variable, say $r_i$, there is only one solution to $\sum_{j=1}^{M}\sum_{k=1}^{Q}\frac{e^{r_i+w_j+c_k}}{e^{r_i+w_j+c_k}+1} = n_i$, since $f(r_i) = \sum_{j=1}^{M}\sum_{k=1}^{Q}\frac{e^{r_i+w_j+c_k}}{e^{r_i+w_j+c_k}+1}$ is a monotonic function in the range [0, N]. This solution can be efficiently found via Newton's method.

---

**Algorithm 1 Iterative strategy to solve Eq. (S4.6)**

**Initialization:** $\{\boldsymbol{r_i}\}$ , $\{\boldsymbol{w_j}\}$**,** and $\{\boldsymbol{u_k}\}$ ← 0s

**While** $\{\boldsymbol{r_i}\}$ , $\{\boldsymbol{w_j}\}$**,** and $\{\boldsymbol{u_k}\}$ are not converged, **do**

// Parallelly update $\{\boldsymbol{r_i}\}$

Parallelly update $\boldsymbol{r_i}$ for $\boldsymbol{i} = \boldsymbol{1}, \dots, \boldsymbol{N}$ by solving $\sum_{j=1}^{M}\sum_{k=1}^{Q}\frac{e^{r_i+w_j+\mu_k}}{e^{r_i+w_j+\mu_k}+1} = \boldsymbol{n_i}$

// Parallelly update $\{\boldsymbol{w_j}\}$

Parallelly update $\boldsymbol{w_j}$ for $\boldsymbol{j} = \boldsymbol{1}, \dots, \boldsymbol{M}$ by solving $\sum_{i=1}^{N}\sum_{k=1}^{Q}\frac{e^{r_i+w_j+\mu_k}}{e^{r_i+w_j+\mu_k}+1} = \boldsymbol{m_j}$

// Parallelly update $\{\boldsymbol{u_k}\}$

Parallelly update $\boldsymbol{\mu_k}$ for $\boldsymbol{k} = \boldsymbol{1}, \dots, \boldsymbol{Q}$ by solving $\sum_{i=1}^{N}\sum_{j=1}^{M}\frac{e^{r_i+w_j+\mu_k}}{e^{r_i+w_j+\mu_k}+1} = \boldsymbol{q_k}$

**end while**

---

4.2 Extensions of CMC Model

4.2.1 Extend to multidimensional binary contingency tensor

Consider a D-dimensional $M_1 \times M_2 \times \cdots \times M_d \times \cdots \times M_D$ contingency tensor. Let $\vec{I}_D = (i_1, i_2, \dots, i_d, \dots, i_D)$ denotes the index of the entry in the D-dimensional tensor, where $i_d$ is the index of the $d^{\text{th}}$ dimension. Let $U_D$ be the set of all possible $\vec{I}_D$s. Define $(i_1, i_2, \dots, i_{d-1}, \cdot_d, i_{d+1}, \dots, i_D)$ as a set of indexes, where "$\cdot_d$" indicates all possible indexes of the $d^{\text{th}}$ dimension (i.e., $\{1, \dots, M_d\}$). Similarly, $(\cdot_1, \dots, \cdot_{d-1}, i_d, \cdot_{d+1}, \dots, \cdot_D)$ is a set of indexes, where only the index of the $d^{\text{th}}$ dimension is fixed to $i_d$ and the indexes of all the other dimensions can be any possible values. $x_{\vec{I}_D}$(=0 or 1) is the observed value in entry $entry_{\vec{I}_D}$ of the tensor. The marginal total of the $i_d^{\text{th}}$ individual in the $d^{\text{th}}$ dimension is $m_{i_d,d} = \sum_{\vec{I}_D \in (\cdot_1, \dots, \cdot_{d-1}, i_d, \cdot_{d+1}, \dots, \cdot_D)} x_{\vec{I}_D}$. For brevity, we will denote $m_{i_d,d}$ as $m_{i_d}$ when there is no confusion.

We assume each entry of the contingency tensor is independent, and follows the Bernoulli distribution with probability $P_{\vec{I}_D}$. $P_{\vec{I}_D}$ can be inferred by maximizing the entropy under the marginal expectation constraints:

$$\max_{\vec{P}} -\sum_{\vec{I}_D \in U_D}\left[P_{\vec{I}_D} \ln P_{\vec{I}_D} + \left(1 - P_{\vec{I}_D}\right)\ln\left(1 - P_{\vec{I}_D}\right)\right]$$
$$s.t. \quad \sum_{\vec{I}_D \in (\cdot_1, \dots, \cdot_{d-1}, i_d, \cdot_{d+1}, \dots, \cdot_D)} P_{\vec{I}_D} = m_{i_d}, \text{ for } i_d = 1, \dots, M_d \text{ and } d = 1, \dots, D \tag{S4.7}$$

To solve Eq. (S4.7), Lagrange multipliers are introduced:

$$\max_{\vec{P}, \vec{r}} -\sum_{\vec{I}_D \in U_D}\left[P_{\vec{I}_D} \ln P_{\vec{I}_D} + \left(1 - P_{\vec{I}_D}\right) ln\left(1 - P_{\vec{I}_D}\right)\right] + \sum_{d=1}^{D} \sum_{i_d=1}^{M_d} r_{i_d}\left(\sum_{\vec{I}_D \in (\cdot_1, \dots, \cdot_{d-1}, i_d, \cdot_{d+1}, \dots, \cdot_D)} P_{\vec{I}_D} - m_{i_d}\right) \tag{S4.8}$$

where $\vec{r} = \left[r_{1,1}, \dots, r_{M_1,1}, r_{1,2}, \dots, r_{M_1,2}, \dots, r_{1,D}, \dots, r_{M_D,D}\right]$. Here $r_{i_d,d}$ is related to the $i_d^{th}$ individual of the $d^{th}$ dimension. Similarly, we will denote $r_{i_d,d}$ as simply $r_{i_d}$ when there is no confusion. $r_{i_d}$ is related to the susceptibility of individual $i_d$ in dimension $d$.

The optimal solution of Eq. (S4.8) is:

$$P_{\vec{I}_D} = \frac{e^{\sum_{d=1}^{D} r_{i_d}}}{e^{\sum_{d=1}^{D} r_{i_d}} + 1} \tag{S4.9}$$

where $\sum_{d=1}^{D} r_{i_d}$ is the sum of susceptivity of individual that the entry $\vec{I}_D$ corresponding to in each of the D dimensions, and $\{r_{i_d}\}$ is the solution to the margin constraints:

$$\sum_{\vec{I}_D \in (\cdot_1, \dots, \cdot_{d-1}, i_d, \cdot_{d+1}, \dots, \cdot_D)} \frac{e^{\sum_{d=1}^{D} r_{i_d}}}{e^{\sum_{d=1}^{D} r_{i_d}} + 1} = m_{i_d} \quad \text{for } i_d = 1, \dots, M_d \text{ and } d = 1, \dots, D \tag{S4.10}$$

4.2.2 Extend to tensor with expected values

For now, the values in the tensor are binary values. However, in practice, we may not be very confident with the observed values. For example, in an experiment observing whether a specific TF binds to a particular gene, according to the experimental data, we conclude that the TF and gene are likely bound together. “Likely” was used because the wrong conclusion might be made due to known/unknown factors, such as measurement error. In such cases, rather than rashly assigning 0 or 1 to the contingency tensor, using an expected value $e$ $(0 \leq e \leq 1)$ is a wise way. In this case, the marginal total would be $m_{i_d} = \sum_{\vec{I}_D \in (\cdot_1,\dots,\cdot_{d-1},i_d,\cdot_{d+1},\dots,\cdot_D)} e_{\vec{I}_D}$, for $i = 1, \dots, M_d$, and $d = 1, \dots, D$. Note that $\{m_{i_d}\}$ may not be integers.

While inferring the probability distribution for such a tensor with expected values, the object function and the margin constraints (Eq. S4.7) are still held in this case. And the solutions are also the same as Eq. (S4.9) and Eq. (S4.10).

4.2.3 Extend to tensor with missing values

Missing values exist in collected datasets of many fields, including science, engineering, and business. In such datasets, only the non-missing (observed) values are used to infer the susceptivities of each factor’s individuals.

For a D-dimensional $M_1 \times M_2 \times \cdots \times M_d \times \cdots \times M_D$ binary tensor, let $\Omega_\mathrm{D}$ denotes the indexes of observed entries of the D-dimension tensor. Assume $(\cdot_1, \dots, \cdot_{d-1}, i_d, \cdot_{d+1}, \dots, \cdot_D) \cap \Omega \neq \emptyset$ for all $i_d$ and all $d$, that is, any individual of any dimension has at least one non-missing value.

To infer the distribution of each entry:

$$\max_{\vec{P}} -\sum_{\vec{I}_D \in \Omega_\mathrm{D}} \left[P_{\vec{I}_D} \ln P_{\vec{I}_D} + \left(1 - P_{\vec{I}_D}\right) \ln\left(1 - P_{\vec{I}_D}\right)\right] \tag{S4.11}$$

$$s.t. \quad \sum_{\vec{I}_D \in (\cdot_1,\dots,\cdot_{d-1},i_d,\cdot_{d+1},\dots,\cdot_D) \cap \Omega_\mathrm{D}} P_{\vec{I}_D} = m_{i_d}, \text{ for } i_d = 1, \dots, M_d \text{ and } d = 1, \dots, D$$

where $\{m_{i_d}\}$ are the marginal totals of the observed values. $m_{i_d} = \sum_{\vec{I}_D \in (\cdot_1,\dots,\cdot_{d-1},i_d,\cdot_{d+1},\dots,\cdot_D) \cap \Omega_\mathrm{D}} x_{\vec{I}_D}$, for $i = 1, \dots, M_d$ and $d = 1, \dots, D$.

The optimal solution is:

$$P_{\vec{I}_D \in \Omega_\mathrm{D}} = \frac{e^{\sum_{d=1}^{D} r_{i_d}}}{e^{\sum_{d=1}^{D} r_{i_d}} + 1} \tag{S4.12}$$

where $\{r_{i_d}\}$ is the solution to the margin constraints:

$$\sum_{\vec{I}_D \in (\cdot_1,\dots,\cdot_{d-1},i_d,\cdot_{d+1},\dots,\cdot_D) \cap \Omega_\mathrm{D}} \frac{e^{\sum_{d=1}^{D} r_{i_d}}}{e^{\sum_{d=1}^{D} r_{i_d}} + 1} = m_{i_d} \quad \text{for } i_d = 1, \dots, M_d;\ d = 1, \dots, D \tag{S4.13}$$

Eq. (S4.12) shares the form with that of tensor without missing values, except that $\{r_{i_d}\}$ are learned from observed entries.

We have learned from Eq. (S4.10) and Eq. (S4.12) that the relationship between the probability distribution of an entry and its corresponding hidden factor $\{r_{i_d}\}$. According to this relationship, the probability distributions of the missing items can also be estimated given that $\{r_{i_d}\}$ has been learned from the observed entries. From this point of view, the CMC model has the potential to impute the missing values in multi-dimensional tensors, especially for datasets that the heterogeneities that exist in multiple dimensions. The imputed values are the expected values of the missing entries given the inferred susceptivities of the corresponding individuals.

#### 4.2.4 Extended to multidimensional integer contingency tensor

We extended the CMC model to handle tensors with integer values, where the integer values follow the binomial distributions with success probabilities $P_{\vec{I}_D}$s to be inferred, given the number of trials $Y_{\vec{I}_D}$s for each entry $\vec{I}_D$.

We first discuss a special case where $Y = Y_{\vec{I}_D}$ for all $\vec{I}_D \in U_D$, then extend to the general cases where $Y_{\vec{I}_D}$s are not identical.

The integer value in each entry is the number of successes in a sequence of $Y$ independent Bernoulli trials, where the trials share an identical success probability. Therefore, the D-dimension integer tensor can be regarded as the sum of $Y$ D-dimension binary tensor, in which each entry of every binary tensor records the outcomes of a single Bernoulli trial. Inspired by this, as shown in Fig. S 14A, for an integer tensor, we introduce a virtual dimension (D+1) to the original D-dimensional tensor. The length of this dimension is equal to $Y$. Along with this virtual dimension, each individual records the results of a single trial in each entry. In this case, we can estimate the probability distribution of this D+1-dimensional binary tensor.

Let's consider a D-dimensional $M_1 \times M_2 \times \cdots \times M_d \times \cdots \times M_D$ integer contingency tensor, in which the value of $\text{entry}_{\vec{I}_D}$ follows the binomial distribution $B(Y, P_{\vec{I}_D})$. A virtual dimension (the (D+1)$^{\text{th}}$ dimension) is introduced, along which each individual is a D-dimensional binary tensor. For the new (D+1)-dimensional $M_1 \times \cdots \times M_D \times M_{D+1}$ binary tensor, $M_{D+1} = Y$, and the value of $\text{entry}_{\vec{I}_{D+1}}$ follows the Bernoulli distribution with success probability $P_{\vec{I}_{D+1}}$. $P_{\vec{I}_{D+1}}$ can be inferred according to the maximum entropy principle, under the marginal expectation constraints:

$$\max_{\vec{P}} \; -\sum\nolimits_{\vec{I}_{D+1}} \left[ P_{\vec{I}_{D+1}} \ln P_{\vec{I}_{D+1}} + \left(1 - P_{\vec{I}_{D+1}}\right) \ln\left(1 - P_{\vec{I}_{D+1}}\right) \right] \qquad \text{(S4.14)}$$

$$s.t. \quad \sum\nolimits_{\vec{I}_{D+1} \in (\cdot_1, \dots, \cdot_{d-1}, i_d, \cdot_{d+1}, \dots, \cdot_{D+1})} P_{\vec{I}_{D+1}} = m_{i_d}, \text{ for } i_d = 1, \dots, M_d;\; d = 1, \dots, D$$

$$P_{(\vec{I}_D, 1_{D+1})} = P_{(\vec{I}_D, i_{D+1})}, \text{ for } i_{D+1} = 2, \dots, M_{D+1};\; \vec{I}_D \in U_D$$

where $(\vec{I}_D,\ i_{D+1})$ denotes the index vector of the (D+1)-dimensional tensor, in which, $\vec{I}_D$ represents the index of the first D dimensions and $i_{D+1}$ denote the index of the last dimension. There is no marginal constrain on the virtual dimension. Instead, an additional constraint set is added to force the entries along the virtual dimension to have the same success probabilities.

To solve Eq. (S4.14), Lagrange multipliers are again introduced:

$$\max_{\vec{P},\vec{r}} -\sum_{\vec{I}_{D+1}}\left[P_{\vec{I}_{D+1}}\ln P_{\vec{I}_{D+1}} + \left(1-P_{\vec{I}_{D+1}}\right)ln\left(1-P_{\vec{I}_{D+1}}\right)\right] + \sum_{d=1}^{D}\sum_{i_d=1}^{M_d} r_{i_d}\left(\sum_{\vec{I}_{D+1}\in(\cdot_1,\dots,\cdot_{d-1},i_d,\cdot_{d+1},\dots,\cdot_{D+1})} P_{\vec{I}_{D+1}} - m_{i_d}\right) + \sum_{\vec{I}_D}\sum_{i_{D+1}=2}^{M_{D+1}}\mu_{(\vec{I}_D,i_{D+1})}\left(P_{(\vec{I}_D,1_{D+1})} - P_{(\vec{I}_D,i_{D+1})}\right) \quad \text{(S4.15)}$$

where $\vec{r} = \{r_{i_d}\}$ for $i = 1,\dots,M_d$ and $d = 1,\dots,D,D+1$.

The optimal solution of Eq. (S4.15) is:

$$P_{(\vec{I}_D,i_{D+1})} = \begin{cases} \frac{e^{\sum_{d=1}^{D} r_{i_d}}}{e^{\sum_{d=1}^{D1} r_{i_d}}+1}, & \text{for } i_{D+1} = 1 \\ \frac{e^{\mu_{(\vec{I}_D,i_{D+1})}+\sum_{d=1}^{D} r_{i_d}}}{e^{\mu_{(\vec{I}_D,i_{D+1})}+\sum_{d=1}^{D} r_{i_d}}+1}, & \text{for } i_{D+1} = 2,\dots,M_{D+1} \end{cases} \quad \text{(S4.16)}$$

where $\{\mu_{(\vec{I}_D,i_{D+1})}\}$ and $\{r_{i_d}\}$ are the solution to:

$$\sum_{\vec{I}_D\in(\cdot_1,\dots,\cdot_{d-1},i_d,\cdot_{d+1},\dots,\cdot_{D+1})}\frac{e^{\sum_{d=1}^{D} r_{i_d}}}{e^{\sum_{d=1}^{D} r_{i_d}}+1} + \sum_{\vec{I}_D\in(\cdot_1,\dots,\cdot_{d-1},i_d,\cdot_{d+1},\dots,\cdot_{D+1})}\sum_{i_{D+1}=2}^{M_{D+1}}\frac{e^{\mu_{(\vec{I}_D,i_{D+1})}+\sum_{d=1}^{D} r_{i_d}}}{e^{\mu_{(\vec{I}_D,i_{D+1})}+\sum_{d=1}^{D} r_{i_d}}+1} = m_{i_d}, \quad \text{for } d = 1,\dots,D \quad \text{(S4.17)}$$

$$\frac{e^{\sum_{d=1}^{D} r_{i_d}}}{e^{\sum_{d=1}^{D} r_{i_d}}+1} = \frac{e^{\mu_{(\vec{I}_D,i_{D+1})}+\sum_{d=1}^{D} r_{i_d}}}{e^{\mu_{(\vec{I}_D,i_{D+1})}+\sum_{d=1}^{D} r_{i_d}}+1}, \text{ for any } \vec{I}_D, \text{ and } i_{D+1} = 2,\dots,M_{D+1} \quad \text{(S4.18)}$$

From Eq. (S4.18), we had $\mu_{(\vec{I}_D,i_{D+1})} = 0$ for any $\vec{I}_D$, and $i_{D+1} = 2,\dots,M_{D+1}$. Therefore, Eq. (S4.16), Eq. (S4.17), and Eq. (S4.18) can be rewritten as:

$$P_{\vec{I}_D} = \frac{e^{\sum_{d=1}^{D} r_{i_d}}}{e^{\sum_{d=1}^{D} r_{i_d}}+1} \quad \text{(S4.19)}$$

$$\sum_{\vec{I}_D\in(\cdot_1,\dots,\cdot_{d-1},i_d,\cdot_{d+1},\dots,\cdot_D)} Y\frac{e^{\sum_{d=1}^{D} r_{i_d}}}{e^{\sum_{d=1}^{D} r_{i_d}}+1} = m_{i_d}, \text{ for } i_d = 1,\dots,M_d \text{ and } d = 1,\dots,D \quad \text{(S4.20)}$$

We extend to the general cases where $Y_{\vec{I}_D}$s are different. For such cases, as shown in Fig. S 14B, we also introduce a virtual dimension (D+1) to the original D-dimensional tensor. The length of this dimension is equal to $Y$, where $Y = \max(Y_{\vec{I}_D})$. For entry$_{\vec{I}_D}$ in the original integer tensor, if its number of trials $Y_{\vec{I}_D}$ is smaller than $Y$, then among its corresponding binary entries, $Y - Y_{\vec{I}_D}$ of them will be marked as missing values. We then estimate the probability distribution of this D+1 dimensional binary tensor with missing values.

Consider a D-dimensional $M_1 \times M_2 \times \cdots \times M_d \times \cdots \times M_D$ integer contingency tensor, in which the value of entry$_{\vec{I}_D}$ follows the binomial distribution $B(Y_{\vec{I}_D}, P_{\vec{I}_D})$. A virtual dimension (the (D+1)$^{\text{th}}$ dimension) is introduced, along which each individual is a D-dimensional binary tensor. For the new $M_1 \times \cdots \times M_D \times M_{D+1}$ binary tensor, $M_{D+1} = Y = \max(Y_{\vec{I}_D})$. For entry$_{\vec{I}_D}$ in the D-dimensional integer tensor, among its corresponding entries in the (D+1)-dimensional binary tensor (entry$_{(\vec{I}_D,1)}, \ldots,$ entry$_{(\vec{I}_D,Y)}$), $Y - Y_{\vec{I}_D}$ of them (entry$_{\left(\vec{I}_D, Y_{\vec{I}_D}+1\right)}, \ldots,$ entry$_{(\vec{I}_D,Y)}$) , are marked as missing values. Let $\Omega_{D+1}$ denotes the indexes of the non-missing entries of the (D+1)-dimensional binary tensor. Value of entry$_{\vec{I}_{D+1}}$ (for $\vec{I}_{D+1} \in \Omega_{D+1}$) follows Bernoulli distribution with success probability $P_{\vec{I}_{D+1}}$. Again, $P_{\vec{I}_{D+1}}$ can be inferred according to the maximum entropy principle, under the marginal expectation constraints:

$$\max_{\vec{P}} \; -\sum_{\vec{I}_{D+1}\in\Omega_{D+1}} \left[P_{\vec{I}_{D+1}} \ln P_{\vec{I}_{D+1}} + \left(1 - P_{\vec{I}_{D+1}}\right) \ln\left(1 - P_{\vec{I}_{D+1}}\right)\right] \quad \text{(S4.21)}$$

$$s.t. \quad \sum_{\vec{I}_{D+1}\in(\cdot_1,\ldots,\cdot_{d-1},i_d,\cdot_{d+1},\ldots,\cdot_{D+1})\cap\Omega_{D+1}} P_{\vec{I}_{D+1}} = m_{i_d}, \text{ for } i_d = 1,\ldots,M_d \text{ and } d = 1,\ldots,D$$

$$P_{(\vec{I}_D,1_{D+1})} = P_{(\vec{I}_D,i_{D+1})}, \text{ for } \left(\vec{I}_D,\ i_{D+1}\right) \in \Omega_{D+1}, \text{ where } \vec{I}_D \in U_D, \text{ and } i_{D+1} = 2,\ldots,Y_{\vec{I}_D}$$

To solve Eq. (S4.21), Lagrange multipliers are again introduced:

$$\begin{gathered}\max_{\vec{P},\vec{r}} - \sum_{\vec{I}_{D+1}\in\Omega_{D+1}} \left[P_{\vec{I}_{D+1}} \ln P_{\vec{I}_{D+1}} + \left(1 - P_{\vec{I}_{D+1}}\right) ln\left(1 - P_{\vec{I}_{D+1}}\right)\right] + \\ \sum_{d=1}^{D}\sum_{i_d=1}^{M_d} r_{i_d}\left(\sum_{\vec{I}_{D+1}\in(\cdot_1,\ldots,\cdot_{d-1},i_d,\cdot_{d+1},\ldots,\cdot_{D+1})\cap\Omega_{D+1}} P_{\vec{I}_{D+1}} - m_{i_d}\right) + \\ \sum_{\vec{I}_D}\sum_{i_{D+1}=2}^{Y_{\vec{I}_D}} \mu_{(\vec{I}_D,i_{D+1})}\left(P_{(\vec{I}_D,1_{D+1})} - P_{(\vec{I}_D,i_{D+1})}\right)\end{gathered} \quad \text{(S4.22)}$$

The optimal solution of Eq. (S4.22) is:

$$P_{(\vec{I}_D,i_{D+1})\in\Omega_{D+1}} = \begin{cases} \dfrac{e^{\sum_{d=1}^{D} r_{i_d}}}{e^{\sum_{d=1}^{D1} r_{i_d}+1}}, & \text{for } i_{D+1} = 1 \\ \dfrac{e^{\mu_{(\vec{I}_D,i_{D+1})}+\sum_{d=1}^{D} r_{i_d}}}{e^{\mu_{(\vec{I}_D,i_{D+1})}+\sum_{d=1}^{D} r_{i_d}}+1}, & \text{for } i_{D+1} = 2,\ldots,M_{D+1} \end{cases} \quad \text{(S4.23)}$$

where $\{\mu_{(\vec{I}_D,i_{D+1})\in\Omega_{D+1}}\}$ and $\{r_{i_d}\}$ are the solution to:

$$\sum_{\vec{I}_D\in(\cdot_1,\ldots,\cdot_{d-1},i_d,\cdot_{d+1},\ldots,\cdot_D)} \frac{e^{\sum_{d=1}^{D} r_{i_d}}}{e^{\sum_{d=1}^{D} r_{i_d}} + 1} + \sum_{\vec{I}_D\in(\cdot_1,\ldots,\cdot_{d-1},i_d,\cdot_{d+1},\ldots,\cdot_D)} \sum_{i_{D+1}=2}^{Y_{\vec{I}_D}} \frac{e^{\mu_{(\vec{I}_D,i_{D+1})}+\sum_{d=1}^{D} r_{i_d}}}{e^{\mu_{(\vec{I}_D,i_{D+1})}+\sum_{d=1}^{D} r_{i_d}}+1} = m_{i_d},$$

$$\text{for } d = 1,\ldots,D \quad \text{(S4.24)}$$

$$\frac{e^{\sum_{d=1}^{D} r_{i_d}}}{e^{\sum_{d=1}^{D} r_{i_d}}+1}=\frac{e^{\mu_{(\vec{I}_D,i_{D+1})}+\sum_{d=1}^{D} r_{i_d}}}{e^{\mu_{(\vec{I}_D,i_{D+1})}+\sum_{d=1}^{D} r_{i_d}}+1}\text{, for } \vec{I}_D \in U_D, \text{and } i_{D+1}=2,\dots,Y_{\vec{I}_D} \quad \text{(S4.25)}$$

From Eq. (S4.25), we have $\mu_{(\vec{I}_D,i_{D+1})\in\Omega_{D+1}}=0$ for $\vec{I}_D \in U_D$, and $i_{D+1}=2,\dots,Y_{\vec{I}_D}$. Therefore, Eq. (S4.23), Eq. (S4.24), and (S4.25) can be rewritten as:

$$P_{\vec{I}_D}=\frac{e^{\sum_{d=1}^{D} r_{i_d}}}{e^{\sum_{d=1}^{D} r_{i_d}}+1} \quad \text{(S4.26)}$$

$$\sum_{\vec{I}_D\in(\cdot_1,\dots,\cdot_{d-1},i_d,\cdot_{d+1},\dots,\cdot_D)} Y_{\vec{I}_D}\frac{e^{\sum_{d=1}^{D} r_{i_d}}}{e^{\sum_{d=1}^{D} r_{i_d}}+1}=m_{i_d}, \quad \text{for } i_d=1,\dots,M_d \text{ and } d=1,\dots,D \quad \text{(S4.27)}$$

Similarly, the CMC model can be extended to integer tensors with missing values. In this case, while expanding the original D-dimensional tensor into (D+1)-dimensional tensor, if the values $\text{entry}_{\vec{I}_D}$ in the original D-dimensional tensor is missing, then all the values of the corresponding entries ($\text{entry}_{(\vec{I}_D,i_{D+1})}$, for all $i_{D+1}$) in the (D+1)-dimensional tensor will also be marked as missing. Similarly, the maximum entropy principle under the marginal expectation constraints and constraints on the $(D+1)^{th}$ dimension is applied, and the success probabilities $P_{\vec{I}_D}$ of each entry in the D-dimensional integer tensor is:

$$P_{\vec{I}_D\in\Omega_D}=\frac{e^{\sum_{d=1}^{D} r_{i_d}}}{e^{\sum_{d=1}^{D} r_{i_d}}+1} \quad \text{(S4.28)}$$

where $\Omega_D$ is the indexes of entiries with non-missing values, and $r_{i_d}$ can be obtained by solving the equations:

$$\sum_{\vec{I}_D\in(\cdot_1,\dots,\cdot_{d-1},i_d,\cdot_{d+1},\dots,\cdot_D)\in\Omega_D} Y_{\vec{I}_D}\frac{e^{\sum_{d=1}^{D} r_{i_d}}}{e^{\sum_{d=1}^{D} r_{i_d}}+1}=m_{i_d}, \quad \text{for } i_d=1,\dots,M_d \text{ and } d=1,\dots,D$$

(S4.29)

Eq. (S4.28) is the optimal distribution for the entry of the *D*+1 dimensional binary tensor. Values in the original *D* dimensional integer tensor are the sum of the *D*+1 dimensional tensor across the virtual dimension. The probability distribution of each entry in the original *D* dimensions tensor is:

$$\Pr\left(x_{\vec{I}_D};Y_{\vec{I}_D},P_{\vec{I}_D}\right)=\binom{Y_{\vec{I}_D}}{x_{\vec{I}_D}}P_{\vec{I}_D}^{x_{\vec{I}_D}}\left(1-P_{\vec{I}_D}\right)^{Y_{\vec{I}_D}-x_{\vec{I}_D}}, \quad \text{for } \vec{I}_D\in\Omega_D \quad \text{(S4.30)}$$

where $P_{\vec{I}_D}$ is given by Eq. (S4.26). When $Y_{\vec{I}_D}$=1 for all $\vec{I}_D \in \Omega_D$, the solutions are the same as those of the binary tensor (Eq. (S4.12) and Eq.(S4.13)).

Similarly to Algorithm 1, Algorithm 2 is proposed to find a solution to Eq. (S4.29), in which, variable sets $\vec{r}_{\cdot d}$ are updated sequentially for $d=1,\dots,D$. And during the update of each variable set $\vec{r}_{\cdot d}$, variable elements within each set are updated parallelly, given the fact that the variable elements within each set are independent if the other variable sets are fixed. For a single variable, say $r_{i_d}$, there is only one solution to $\sum_{\vec{I}_D \in (\cdot_1,\ldots,\cdot_{d-1}, i_d, \cdot_{d+1},\ldots,\cdot_D) \in \Omega_D} Y_{\vec{I}_D} \frac{e^{\sum_{d=1}^{D} r_{i_d}}}{e^{\sum_{d=1}^{D} r_{i_d}} + 1} = m_{i_d}$, since $(r_{i_d}) = \sum_{\vec{I}_D \in (\cdot_1,\ldots,\cdot_{d-1}, i_d, \cdot_{d+1},\ldots,\cdot_D) \in \Omega_D} Y_{\vec{I}_D} \frac{e^{\sum_{d=1}^{D} r_{i_d}}}{e^{\sum_{d=1}^{D} r_{i_d}} + 1}$ is a monotonic function. This solution can be efficiently found via Newton's method. Algorithm 2 is guaranteed to find a solution to Eq. (S4.29) (Theorem 4.3).

**Algorithm 2 Iterative strategy to solve Eq. (S4.29)**

**Initialization:** $\vec{r} \leftarrow$ 0s
**while** $\vec{r}$ is not converged, **do**
  **for** $d$ = 1 **to** $D$  // for each dimension
    Parallelly update $r_{i_d}$ for $i_d = 1, \ldots, M_d$ by solving $f(r_{i_d}) = m_{i_d}$
    using Newton's method
  **end**
**end while**

4.3 Properties of CMC Model

In this section, we will prove three CMC properties that we mentioned before:

**Property1**: The probability distribution defined by Eq. (S4.28) and Eq. (S4.29) is unique. This property will be formally stated and proved in **Theorem 4.1.**

**Property2**: The ratio between any two variables of the same factors, $\frac{e^{r_{i_d}}}{e^{r_{i'_d}}}$, is fixed for $i_d, i'_d = 1, \ldots, M_d$ and $d = 1, \ldots, D$ (**Theorem 4.2**).

**Property3**: The iterative update of $\vec{r}$ (Algorithm 2) will converge to a solution that satisfies Equation (S4.29) (**Theorem 4.3**).

First, we will give some definitions.

For a D dimensional $M_1 \times M_2 \times \cdots \times M_d \times \cdots \times M_D$ tensor, define index of entry $\vec{I}_D = (i_1, i_2, \ldots, i_d, \ldots, i_D)$, $i_d = 1, \ldots, M_d$, for $d = 1, \ldots, D$; $i_d$ is the index of the $d^{\text{th}}$ dimension.

Define $\Omega$ as the indexes of items with non-missing values and $(\cdot, \ldots, \cdot, i_d, \cdot, \ldots, \cdot) \cap \Omega \neq \emptyset$.

Define $\vec{I}_D$ induced set $\mathcal{B}(\vec{I}_D)$ as $\{1\} \cup \{d | i_d < M_d\}$; in other words, we pick the first dimension, as well as all other dimensions $d$ whose corresponding coordinate $i_d$ is less than $M_d$.

Let $\vec{r}' = [r'_{1,1}, \dots, r'_{M_1,1}, r'_{1,2}, \dots, r'_{M_2-1,2}, \dots, r'_{1,D}, \dots, r'_{M_D-1,D}]$, and $r'_{i_d,d}$ is shortened as $r'_{i_d}$ when there is no confusion.

We define $m_{i_d,d}$ as the marginal total for the $i_d$th item in dimension $d$, and we also abbreviate it as $m_{i_d}$. These values are calculated from data, and are considered as fixed.

Before we prove Theorem 4.1, we need two lemmas.

**Lemma 4.1** Let

$$f(\vec{r}') = \sum_{\vec{I}_D \in \Omega} Y_{\vec{I}_D} \ln(e^{\sum_{d \in B(\vec{I}_D)} r'_{i_d}} + 1) - (\sum_{i_1=1}^{M_1} r'_{i_1} m_{i_1} + \sum_{d=2}^{D} \sum_{i_d=1}^{M_d} r'_{i_d} m_{i_d})$$

Then $f(\vec{r}')$ is a strictly convex function. Here $Y_{\vec{I}_D}$s are constant values.

*Proof:*

The Hessian matrix of $f(\vec{r}')$

$$H(\vec{r}') = \begin{bmatrix} A_{11}^{M_1 \times M_1} & A_{12}^{M_1 \times (M_2-1)} & \dots & A_{1d}^{M_1 \times (M_d-1)} & \dots & A_{1D}^{M_1 \times (M_D-1)} \\ A_{21}^{(M_2-1) \times M_1} & A_{22}^{(M_2-1) \times (M_2-1)} & \dots & A_{2d}^{(M_2-1) \times (M_d-1)} & \dots & A_{2D}^{(M_2-1) \times (M_D-1)} \\ \vdots & \vdots & \ddots & \vdots & \ddots & \vdots \\ A_{d1}^{(M_d-1) \times M_1} & A_{d2}^{(M_d-1) \times (M_2-1)} & \dots & A_{dd}^{(M_d-1) \times (M_d-1)} & \dots & A_{dD}^{(M_d-1) \times (M_D-1)} \\ \vdots & \vdots & \ddots & \vdots & \ddots & \vdots \\ A_{D1}^{(M_D-1) \times M_1} & A_{D2}^{(M_D-1) \times (M_2-1)} & \dots & A_{Dd}^{(M_D-1) \times (M_d-1)} & \dots & A_{DD}^{(M_D-1) \times (M_D-1)} \end{bmatrix},$$

Each element of $H(\vec{r}')$ is the second order derivative of $f(\vec{r}')$ for each pair of $r'_{i_d}$ ($i = 1, \dots, M_d$ if $d = 1$, and $i = 1, \dots, M_d - 1$ if $d = 2, \dots, D$). $H(\vec{r}')$ is a symmetric matrix, where $A_{dd'} = A_{d'd}^T$ for $d = 1, \dots, D$.

Submatrices $A_{dd}$s are diagonal matrices.

The diagonal elements in $A_{11}^{M_1 \times M_1}$ are:

$$a_{i_1 i_1} = \sum_{\vec{I}_D \in (i_1, \cdot_2, \dots, \cdot_D) \cap \Omega} Y_{\vec{I}_D} \frac{e^{\sum_{d' \in \mathcal{B}(\vec{I}_D)} r'_{i_{d'}}}}{\left(e^{\sum_{d' \in \mathcal{B}(\vec{I}_D)} r'_{i_{d'}}} + 1\right)^2}, \text{ for } i_1 = 1, \dots, M_1$$

The diagonal elements in $A_{dd}^{(M_d-1) \times (M_d-1)}$ (where d>1) are:

$$a_{i_d i_d} = \sum_{\vec{I}_D \in (\cdot_1, \dots, \cdot_{d-1}, i_d, \cdot_{d+1}, \dots, \cdot) \cap \Omega} Y_{\vec{I}_D} \frac{e^{\sum_{d' \in \mathcal{B}(\vec{I}_D)} r'_{i_{d'}}}}{\left(e^{\sum_{d' \in \mathcal{B}(\vec{I}_D)} r'_{i_{d'}}} + 1\right)^2}, \text{ for } i_d = 1, \dots, M_d - 1$$

For $A_{1d}^{M_1 \times (M_d-1)}$ where $d = 2, \dots, D$, its elements are:

$$a_{i_1 i_d} = \sum_{\vec{I}_D \in (i_1, \cdot_2, \dots, \cdot_{d-1}, i_d, \cdot_{d+1}, \dots, \cdot_D) \cap \Omega} Y_{\vec{I}_D} \frac{e^{\sum_{d' \in \mathcal{B}(\vec{I}_D)} r'_{i_{d'}}}}{\left(e^{\sum_{d' \in \mathcal{B}(\vec{I}_D)} r'_{i_{d'}}} + 1\right)^2},$$

for $i_1 = 1, \dots, M_1$ and $i_d = 1, \dots, M_d - 1$ if $d > 1$.

For $A_{dd'}$ where $d \neq d'$, $d > 1$, and $d' > 1$, its elements are:

$$a_{i_d i_{d'}} = \sum_{\vec{I}_D \in (\cdot, \dots, \cdot, i_d, \cdot, \dots, \cdot, i_{d'}, \cdot, \dots, \cdot) \cap \Omega} Y_{\vec{I}_D} \frac{e^{\sum_{d'' \in \mathcal{B}(\vec{I}_D)} r'_{i_{d''}}}}{\left(e^{\sum_{d'' \in \mathcal{B}(\vec{I}_D)} r'_{i_{d''}}} + 1\right)^2}, \text{ for } i_d = 1, \dots, M_d - 1.$$

where $(\cdot, \dots, \cdot, i_d, \cdot, \dots, \cdot, i_{d'}, \cdot, \dots, \cdot)$ is shorten for $(\cdot_1, \dots, \cdot_{d-1}, i_d, \cdot_{d+1}, \dots, \cdot_{d'-1}, i_{d'}, \cdot_{d'+1}, \dots, \cdot_D)$. Since $(\cdot_1, \dots, \cdot_{d-1}, i_d, \cdot_{d+1}, \dots, \cdot_D) \cap \Omega \neq \emptyset$, all entries in $H(\vec{r}')$ are non-zeros except the off-diagonal entries in the submatrices $A_{dd}$s.

Let $\vec{x} \in \mathbb{R}^{M_1 + \sum_{d=2}^{D}(M_d - 1)} \backslash \{0\}$, then

$$\vec{x}^T H \vec{x} = \sum_{\vec{I}_D \in \Omega} \left[ Y_{\vec{I}_D} \frac{e^{\sum_{d' \in \mathcal{B}(\vec{I}_D)} r'_{i_{d'}}}}{\left(e^{\sum_{d' \in \mathcal{B}(\vec{I}_D)} r'_{i_{d'}}} + 1\right)^2} \left(y_{i_1} + \sum_{d=2}^{D} y_{i_d}\right)^2 \right],$$

where $y_{i_d} = \begin{cases} 0, & if\ i_d = M_d\ and\ d > 1 \\ x_{i_d}, & \text{others} \end{cases}$. It's obvious that $\vec{x}^T H \vec{x} > 0$, and Hessian matrix $H$ is positive-definite. Therefore, $f(\vec{r}')$ is strictly convex. ■

**Lemma 4.2** Define equations

$$\begin{cases} \displaystyle\sum_{\vec{I}_D \in (i_1, \cdot_2, \dots, \cdot_D) \cap \Omega} Y_{\vec{I}_D} \frac{e^{\sum_{d \in \mathcal{B}(\vec{I}_D)} r'_{i_d}}}{e^{\sum_{d \in \mathcal{B}(\vec{I}_D)} r'_{i_d}} + 1} = m_{i_1}, \text{ for } i_1 = 1, \dots, M_1 \\ \displaystyle\sum_{\vec{I}_D \in (\cdot_1, \dots, \cdot_{d-1}, i_d, \cdot_{d+1}, \dots, \cdot_D)} Y_{\vec{I}_D} \frac{e^{\sum_{d \in \mathcal{B}(\vec{I}_D)} r'_{i_d}}}{e^{\sum_{d \in \mathcal{B}(\vec{I}_D)} r'_{i_d}} + 1} = m_{i_d}, \text{ for } i_d = 1, \dots, M_d - 1 \text{ and } d = 2, \dots, D \end{cases}$$

$Y_{\vec{I}_D}$ are constant values, and $0 < m_{i_d} < \sum_{\vec{I}_D \in (\cdot_1, \dots, \cdot_{d-1}, i_d, \cdot_{d+1}, \dots, \cdot_D) \cap \Omega} Y_{\vec{I}_D}$ for $i = 1, \dots, M_d$ and $d = 1, \dots, D$,. Then the equation set has at most one solution.

*Proof:*

Define $f(\vec{r}') = \sum_{\vec{I}_D \in \Omega} Y_{\vec{I}_D} \ln(e^{\sum_{d \in \{1, d | i_d < M_d\}} r'_{i_d}} + 1) - (\sum_{i_1=1}^{M_1} r'_{i_1} m_{i_1} + \sum_{d=2}^{D} \sum_{i_d=1}^{M_d - 1} r'_{i_d} m_{i_d})$,

where $\vec{r}' = \left[r'_{1_1}, \dots, r'_{M_1 1}, r'_{1_2}, \dots, r'_{(M_2-1)_2}, \dots, r'_{1_D}, \dots, r'_{(M_D-1)_D}\right]$.

From **Lemma 4.1**, we know that $f(\vec{r}')$ is strictly convex. Therefore, any local minimum of $f(\vec{k},\vec{q})$ is also the unique global minimum of $f(\vec{k},\vec{q})$.

To obtain the unique global optimal solution, we take the derivative with respect to each element of $\vec{r}'$ respectively and set them equal to 0:

$$\begin{cases} \sum_{\vec{I}_D \in (i_1, \cdot_2, \dots, \cdot_D) \cap \Omega} Y_{\vec{I}_D} \dfrac{e^{\sum_{d \in \mathcal{B}(\vec{I}_D)} r'_{i_d}}}{e^{\sum_{d \in \mathcal{B}(\vec{I}_D)} r'_{i_d}} + 1} = m_{i_1}, \quad \text{for } i_1 = 1, \dots, M_1 \\ \sum_{\vec{I}_D \in (\cdot_1, \dots, \cdot_{d-1}, i_d, \cdot_{d+1}, \dots, \cdot_D) \cap \Omega} Y_{\vec{I}_D} \dfrac{e^{\sum_{d \in \mathcal{B}(\vec{I}_D)} r'_{i_d}}}{e^{\sum_{d \in \mathcal{B}(\vec{I}_D)} r'_{i_d}} + 1} = m_{i_d}, \quad \text{for } i_d = 1, \dots, M_d - 1 \text{ and } d = 2, \dots, D \end{cases}$$

In other words, any solutions to the equation set are the optimal solutions to $f(\vec{r}')$. Since $f(\vec{r}')$ has at most one global minimum, the equation set has at most one solution.

To prove the equation set has exactly one solution, in the following, we prove that $f(\vec{r}')$ will not obtain its optimal solution when $r'_{i_d} \to \infty$ or $r'_{i_d} \to -\infty$:

For any $r'_{i_d} \in \vec{r}'$, if $r'_{i_d} \to \infty$, then

$$\frac{\partial f}{\partial r'_{i_d}} = \sum_{\vec{I}_D \in (\cdot_1, \dots, \cdot_{d-1}, i_d, \cdot_{d+1}, \dots, \cdot_D) \cap \Omega} Y_{\vec{I}_D} \frac{e^{\sum_{d \in \mathcal{B}(\vec{I}_D)} r'_{i_d}}}{e^{\sum_{d \in \mathcal{B}(\vec{I}_D)} r'_{i_d}} + 1} - m_{i_d}$$
$$= \sum_{\vec{I}_D \in (\cdot_1, \dots, \cdot_{d-1}, i_d, \cdot_{d+1}, \dots, \cdot_D) \cap \Omega} Y_{\vec{I}_D} - m_{i_d}$$

As $m_{i_d} < \sum_{\vec{I}_D \in (\cdot_1, \dots, \cdot_{d-1}, i_d, \cdot_{d+1}, \dots, \cdot_D) \cap \Omega} Y_{\vec{I}_D}$, $\frac{\partial f}{\partial r'_{i_d}} > 0$.

Similarly, if $r'_{i_d} \to -\infty$, then

$$\frac{\partial f}{\partial r'_{i_d}} = \sum_{\vec{I}_D \in (\cdot_1, \dots, \cdot_{d-1}, i_d, \cdot_{d+1}, \dots, \cdot_D) \cap \Omega} Y_{\vec{I}_D} \frac{e^{\sum_{d \in \mathcal{B}(\vec{I}_D)} r'_{i_d}}}{e^{\sum_{d \in \mathcal{B}(\vec{I}_D)} r'_{i_d}} + 1} - m_{i_d}$$
$$= 0 - m_{i_d}$$

As $m_{i_d} > 0$, $\frac{\partial f}{\partial r'_{i_d}} < 0$.

To summarize, $f(\vec{r}')$ will obtain an optimal solution with finite $r'_{i_d}$. Therefore, $f(\vec{r}')$ has and only has one optimal global solution, and the equation set has and only has one solution. ■

**Theorem 4.1** For a D-dimensional $M_1 \times M_2 \times \cdots \times M_d \times \cdots \times M_D$ tensor. Let $\vec{I}_D = (i_1, i_2, \dots, i_d, \dots, i_D)$ denotes the index of the entry in the D-dimensional tensor, where $i_d$ is the index of the $d^{th}$ dimension. Define $(i_1, i_2, \dots, i_{d-1}, \cdot, i_{d+1}, \dots, i_D)$ as a set of indexes, where "·" indicates all possible indexes of the $d^{th}$ dimension (i.e., $i_d \in \{1, \dots, M_d\}$). Similarly, $(\cdot, \dots, \cdot, i_d, \cdot, \dots, \cdot)$ is a set of indexes, where only the index of the $d^{th}$ dimension is fixed to $i_d$ and the indexes of all the other dimensions can be any possible values. Define the probability distribution of each entry:

$$P_{\vec{I}_D} = \frac{e^{\sum_{d=1}^{D} r_{i_d}}}{e^{\sum_{d=1}^{D} r_{i_d}} + 1}$$

where $\{r_{i_d}\}$ is the solution to the constraints:

$$\sum_{\vec{I}_D \in (\cdot_1, \dots, \cdot_{d-1}, i_d, \cdot_{d+1}, \dots, \cdot_D) \cap \Omega} Y_{\vec{I}_D} \frac{e^{\sum_{d=1}^{D} r_{i_d}}}{e^{\sum_{d=1}^{D} r_{i_d}} + 1} = m_{i_d} \quad \text{for } i_d = 1, \dots, M_d; d = 1, \dots, D$$

where $\{m_{i_d}\}$ are the marginal totals and are given, $0 < m_{i_d} < \sum_{\vec{I}_D \in (\cdot_1, \dots, \cdot_{d-1}, i_d, \cdot_{d+1}, \dots, \cdot_D) \cap \Omega} Y_{\vec{I}_D}$ for $i_d = 1, \dots, M_d$ and $d = 1, \dots, D$, and $\Omega$ reprents the indexes of items with non-missing values and $(\cdot_1, \dots, \cdot_{d-1}, i_d, \cdot_{d+1}, \dots, \cdot_D) \cap \Omega \neq \emptyset$. Then among all the solutions $\{r_{i_d}\}$s, $\sum_{d=1}^{D} r_{i_d}$ is always fixed for any $\vec{I}_D$, and $P_{\vec{I}_D}$ has a unique solution.

*Proof:*

Define

$$\begin{cases} r'_{i_1} = r_{i_1} + \sum_{d=2}^{D} r_{M_d} \\ r'_{i_2} = r_{i_2} - r_{M_2} \\ \vdots \\ r'_{i_D} = r_{i_D} - r_{M_D} \end{cases}, \text{ for } i_d = 1, \dots, M_d, \text{ and } d = 1, \dots, D.$$

Then, $r'_{i_d} = 0$, for $i_d = M_d$ and $d = 2, \dots, D$. And $\sum_{d=1}^{D} r_{i_d} = \sum_{d=1}^{D} r'_{i_d}$, for $i_d = 1, \dots, M_d$, and $d = 1, \dots, D$.

The probability function and constraints can be rewritten as:

$$P_{\vec{I}_D} = \frac{e^{\sum_{d=1}^{D} r'_{i_d}}}{e^{\sum_{d=1}^{D} r'_{i_d}} + 1}$$

and

$$\sum_{\vec{I}_D \in (\cdot_1, \dots, \cdot_{d-1}, i_d, \cdot_{d+1}, \dots, \cdot_D) \cap \Omega} Y_{\vec{I}_D} \frac{e^{\sum_{d=1}^{D} r'_{i_d}}}{e^{\sum_{d=1}^{D} r'_{i_d}} + 1} = m_{i_d}, \text{ for } i_d = 1, \dots, M_d, \text{ and } d = 1, \dots, D.$$

As $r'_{i_d}$s are only involved in the addition operation and $r'_{M_d,d} = 0$ for $d = 2, \dots, D$, variables $r'_{M_d,d}$ (for $d = 2, \dots, D$) can be safely omitted during the following procedures. Besides, the constraints $\sum_{\vec{I}_D\in(\cdot_1,\dots,\cdot_{d-1},M_d,\cdot_{d+1},\dots,\cdot_D)} \frac{e^{\sum_{d=1}^{D} r'_{i_d}}}{e^{\sum_{d=1}^{D} r'_{i_d}}+1} = m_{M_d,d}$ for $d = 2,\dots,D$ are redundant and can also be omitted because the sums of marginal totals in different dimension are always the same, i.e., $\sum_{i_1=1}^{M_1} m_{i_1} = \sum_{i_2=1}^{M_2} m_{i_2} = \cdots = \sum_{i_D=1}^{M_D} m_{i_D}$.

Therefore, the constraint equation set can be rewriting as:

$$\begin{cases} \displaystyle\sum_{\vec{I}_D\in(i_1,\cdot_2,\dots,\cdot_D)\cap\Omega} Y_{\vec{I}_D} \frac{e^{\sum_{d\in\mathcal{B}(\vec{I}_D)} r'_{i_d}}}{e^{\sum_{d\in\mathcal{B}(\vec{I}_D)} r'_{i_d}}+1} = m_{i_1}, \text{ for } i_1 = 1,\dots,M_1 \\ \displaystyle\sum_{\vec{I}_D\in(\cdot_1,\dots,\cdot_{d-1},i_d,\cdot_{d+1},\dots,\cdot_D)\cap\Omega} Y_{\vec{I}_D} \frac{e^{\sum_{d\in\mathcal{B}(\vec{I}_D)} r'_{i_d}}}{e^{\sum_{d\in\mathcal{B}(\vec{I}_D)} r'_{i_d}}+1} = m_{i_d}, \text{ for } i_d = 1,\dots,M_d-1 \text{ and } d = 2,\dots,D \end{cases}$$

From **Theorem 4.1**, we know that the constraint equation has a unique solution $\{{r'_{i_d}}^*\}$.

Therefore, $P_{\vec{I}_D} = \frac{e^{\sum_{d=1}^{D} r'_{i_d}}}{e^{\sum_{d=1}^{D} r'_{i_d}+1}}$ is unique, and $\sum_{d=1}^{D} r_{i_d} = \sum_{d=1}^{D} r'_{i_d}$ is fixed. ∎

**Theorem 4.2** If $e^{\sum_{d=1}^{D} r_{i_d}}$ is fixed for $i_d = 1,\dots,M_d$ and $d = 1,\dots,D$, then $\frac{e^{r_{i_d}}}{e^{r_{i'_d}}}$ is fixed for $i_d = 1,\dots,M_d$, $i'_d = 1,\dots,M_d$ and $d = 1,\dots,D$.

*Proof:*

As $e^{\sum_{d=1}^{D} r_{i_d}}$ is fixed, $\sum_{d=1}^{D} r_{i_d}$ is also fixed. Let $\sum_{d=1}^{D} r_{i_d} = c_{I_D}$, where $c_{I_D}$ is a constant value for $I_D = (i_1, i_2, \dots, i_d, \dots, i_D)$.

For any $d^* \in \{1,\dots,D\}$, $r_{i_{d^*}} - r_{i'_{d^*}} = \sum_{d=1}^{d^*-1} r_{i_d} + r_{i_{d^*}} + \sum_{d=d^*+1}^{D} r_{i_d} - \left(\sum_{d=1}^{d^*-1} r_{i_d} + r_{i'_{d^*}} + \sum_{d=d^*+1}^{D} r_{i_d}\right) = c_{(i_1,i_2,\dots,i_{d^*},\dots,i_D)} - c_{(i_1,i_2,\dots,i'_{d^*},\dots,i_D)}$, which is a constant for any $i_{d^*} \in \{1,\dots,M_{d^*}\}$, and $i'_{d^*} \in \{1,\dots,M_{d^*}\}$.

Therefore, $\frac{e^{r_{i_d}}}{e^{r_{i'_d}}}$ is fixed for $i_d = 1,\dots,M_d$, $i'_d = 1,\dots,M_d$ and $d = 1,\dots,D$. ∎

Next, we will prove the convergence of the updating algorithm. We will first introduce a useful lemma.

**Lemma 4.3** (49) Given a real-valued continuous function $h$ on $X \times Y$, define a point-to-set map $\mathcal{A}: X \rightarrow Y$ with respect to $h$ as

$$\mathcal{A}(x) = arg \min_{y' \in Y} h(x, y') = \{y: h(x, y) \leq h(x, y'), \forall y' \in Y\}.$$

If $\mathcal{A}(x)$ is nonempty, $\mathcal{A}$ is closed at $x$.

**Theorem 4.3** The iteratively update operation in Algorithm 2 will converge to a solution that satisfying the constraint equation set Eq. (S4.29).

Let $\mathcal{A}: R \rightarrow \mathcal{P}(R)$ be the point to set map defined by Algorithm 2, where $R$ is a subset of $\mathbb{R}^n$, with $n = \sum_{d=1}^{D} M_d$. Repeated application of $\mathcal{A}$ generates a sequence $\{\vec{r}^{(k)}\}_{k=0}^{\infty}$ for the given initial point $\vec{r}^{(0)} \in R$ through the iteration $\vec{r}^{(k+1)} \in \mathcal{A}(\vec{r}^{(k)})$, where $\vec{r} = [\vec{r}_{\cdot_1}, \dots, \vec{r}_{\cdot_d}, \dots, \vec{r}_{\cdot_D}]$, $\vec{r}_{\cdot_d} = [r_{1,d}, \dots, r_{M_d,d}]$ for $d = 1, \dots, D,$ and $\vec{r}^{(0)} = [0, \dots, 0]$. Let $\Gamma$ be the solution set.

Define continuous function $f(\vec{r}) = \sum_{\vec{I}_D \in \Omega} Y_{\vec{I}_D} \ln(e^{\sum_d r_{i_d}} + 1) - (\sum_{d=1}^{D} \sum_{i_d=1}^{M_d} r_{i_d} m_{i_d})$. Taking derivative w.r.t. $r_{i_d}$ for all $i_d$ gives us:

$$\sum_{\vec{I}_D \in (\cdot_1, \dots, \cdot_{d-1}, i_d, \cdot_{d+1}, \dots, \cdot_D) \cap \Omega} Y_{\vec{I}_D} \frac{e^{\sum_{d=1}^{D} r_{i_d}}}{e^{\sum_{d=1}^{D} r_{i_d}} + 1} = m_{i_d}, \quad \text{for } i_d = [1, \dots, M_d] \text{ and } d = [1, \dots, D]$$

which is exactly the equation set Algorithm 2 to be solved. In the $k+1$ iteration ($k = 0,1,2, \dots$), $\vec{r}_{\cdot_d}^{(k+1)}$ are updated for $d = 1, \dots, D$ one after another. For $d = d'$ (where $d' = 1, \dots, D$), given the values of $\sum_{d=1}^{d'-1} r_{i_d}^{(k+1)}$ and $\sum_{d=d'+1}^{D} r_{i_d}^{(k)}$, $\vec{r}_{\cdot_{d'}}^{(k+1)}$ is computed by solving the equation set

$$\sum_{\vec{I}_D \in (\cdot_1, \dots, \cdot_{d-1}, i'_d, \cdot_{d+1}, \dots, \cdot_D) \cap \Omega} Y_{\vec{I}_D} \frac{e^{\sum_{d=1}^{d'-1} r_{i_d}^{(k+1)} + r_{i_{d'}}^{(k+1)} + \sum_{d=d'+1}^{D} r_{i_d}^{(k)}}}{e^{\sum_{d=1}^{d'-1} r_{i_d}^{(k+1)} + r_{i_{d'}}^{(k+1)} + \sum_{d=d'+1}^{D} r_{i_d}^{(k)}} + 1} = m_{i_{d'}} \quad \text{for } i_{d'} = 1, \dots, M_{d'}.$$

Therefore,

$$f\left(\vec{r}_{i_1}^{(k+1)}, \dots, \vec{r}_{i_{d'-1}}^{(k+1)}, \vec{r}_{i_{d'}}^{(k+1)}, \vec{r}_{i_{d'+1}}^{(k)}, \dots, \vec{r}_{i_D}^{(k)}\right) \leq f\left(\vec{r}_{i_1}^{(k+1)}, \dots, \vec{r}_{i_{d'-1}}^{(k+1)}, \vec{r}_{i_{d'}}^{(k)}, \vec{r}_{i_{d'+1}}^{(k)}, \dots, \vec{r}_{i_D}^{(k)}\right).$$

After computing $\vec{r}_{\cdot_{d'}}^{(k+1)}$ for all $d'$, we have

$$f(\vec{r}^{(k+1)}) \leq f(\vec{r}^{(k)}).$$

If $\vec{r}^{(k)} \in \Gamma$, then the equality holds. If $\vec{r}^{(k)} \notin \Gamma$, then $f(\vec{r}^{(k+1)}) < f(\vec{r}^{(k)})$.

Besides, define a continuous function $h(\vec{x},\vec{y}) = \sum_{d=1}^{D} a^{D-d} f(\vec{y}_{\cdot_1}, \dots, \vec{y}_{\cdot_d}, \vec{x}_{\cdot_{d+1}}, \dots, \vec{x}_{\cdot_D})$ where $a > 0$ is a constant, $\vec{x} \in \mathbb{R}^n$, and $\vec{y} \in \mathbb{R}^n$. Then $\mathcal{A}(\vec{r}) = arg \min_{y' \in \mathbb{R}^n} h(\vec{r},\vec{y}') = \{\vec{y}: h(\vec{r},\vec{y}) \leq h(\vec{r},\vec{y}'), \forall \vec{y}' \in \mathbb{R}^n\}$ as $a \to \infty$. $\mathcal{A}(\vec{r})$ is nonempty for every $\vec{r} \in R$. According to Lemma 4.3, we obtain that $\mathcal{A}(\vec{r})$ is closed on $R$.

In the following, we will prove that all point $\vec{r}^{(k)}$ are in a compact set $\mathcal{S} \subset R$:

Again, $r_{i_{d'}}^{(k+1)}$ is updated by solving:

$$\sum_{\vec{I}_D \in (\cdot_1, \dots, \cdot_{d-1}, i'_d, \cdot_{d+1}, \dots, \cdot_D) \cap \Omega} Y_{\vec{I}_D} \frac{e^{\sum_{d=1}^{d'-1} r_{i_d}^{(k+1)} + r_{i_{d'}}^{(k+1)} + \sum_{d=d'+1}^{D} r_{i_d}^{(k)}}}{e^{\sum_{d=1}^{d'-1} r_{i_d}^{(k+1)} + r_{i_{d'}}^{(k+1)} + \sum_{d=d'+1}^{D} r_{i_d}^{(k)}} + 1} = m_{i_{d'}}$$

Since $0 < m_{i_{d'}} < \sum_{\vec{I}_D \in (\cdot_1, \dots, \cdot_{d-1}, i'_d, \cdot_{d+1}, \dots, \cdot_D) \cap \Omega} Y_{\vec{I}_D}$, the values of $\sum_{d=1}^{d'-1} r_{i_d}^{(k+1)} + r_{i_{d'}}^{(k+1)} + \sum_{d=d'+1}^{D} r_{i_d}^{(k)}$ that solve the equation is finite. If all the elements of $\vec{r}^{(k)}$ are finite, then $r_{i_1}^{(k+1)}$ is finite, and consequently, $r_{i_{d'}}^{(k+1)}$ is finite for all $d'$. In another word, there exists a large finite positive value $b$ such that $r_{i_{d'}}^{(k+1)} \in [-b, b]$ for all $d'$. Therefore, $\vec{r}^{(k+1)}$ is in a compact set $\mathcal{S}^{(k+1)} \subset R$.

As the given initial point $\vec{r}^{(0)} = [0, \dots, 0]$ does not include any infinite values, it can be deduced that $\vec{r}^{(k)}$ for $k = 0,1,2, \dots, \infty$ are in a compact set $\mathcal{S} \subset R$.

To summarize, 1) all point $\vec{r}^{(k)}$ are in a compact set; 2) $\mathcal{A}(\vec{r})$ is closed at $\vec{r} \in R$; 3) and there is a continuous function $f(\vec{r})$, such that: $f(\vec{r}^{(k+1)}) < f(\vec{r}^{(k)})$ if $\vec{r}^{(k)} \notin \Gamma$, and $f(\vec{r}^{(k+1)}) = f(\vec{r}^{(k)})$ if $\vec{r}^{(k)} \in \Gamma$. According to Zangwill's global convergence theorem (50), the limit of $\{\vec{r}^{(k)}\}$ is in $\Gamma$.

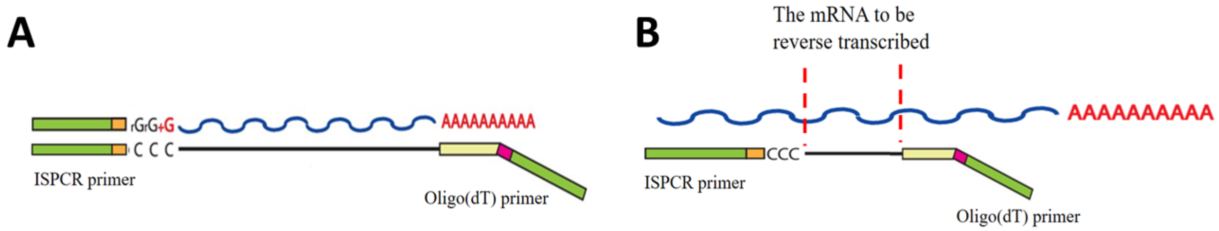


**Fig. S 1.** (A) Reverse transcription (normal case); (B) Incomplete reverse transcription results in truncated cDNA.

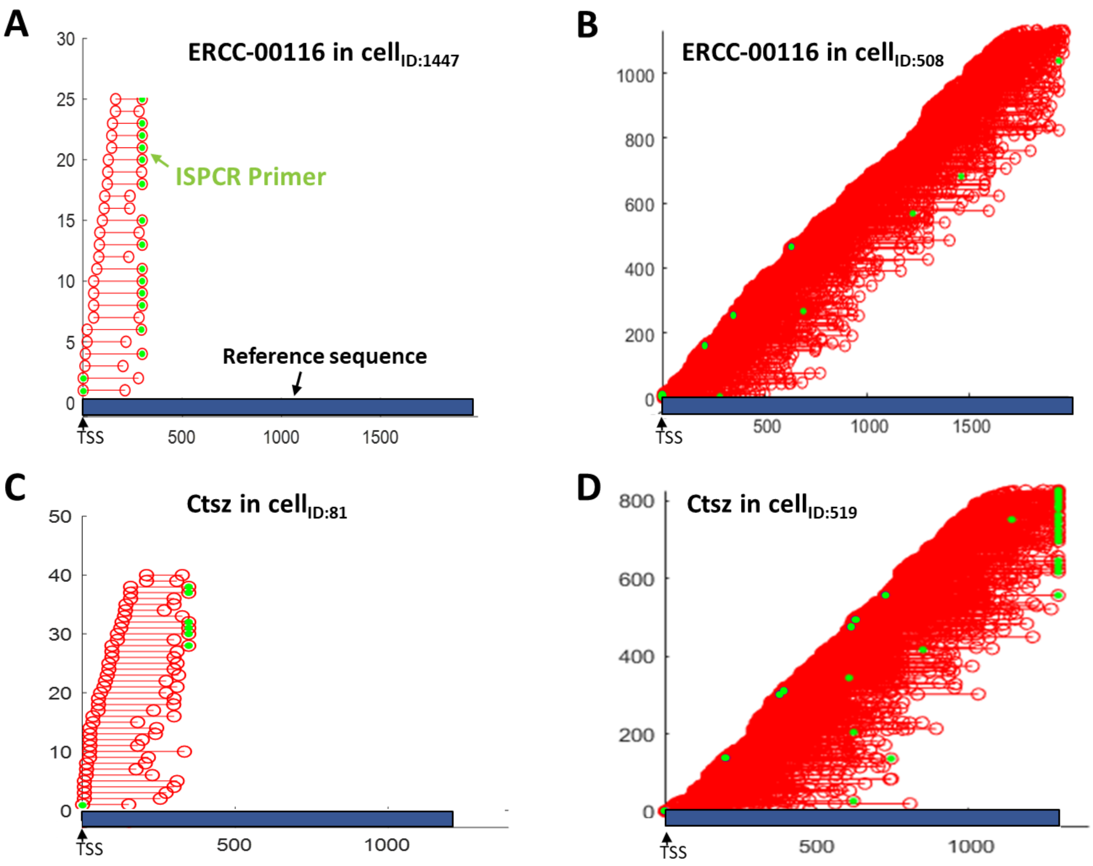


**Fig. S 2**. Results of aligning the sequenced fragments to the reference sequences (introns are omitted). In the figure, one red line with two red circles in both ends represents a sequenced fragment. The horizontal axis represents the position where the sequenced fragments align to. All the sequence reads are ordered according to their starting points of the alignment positions. Greed circles indicate ISPCR primers are found in the corresponding fragments' ends.

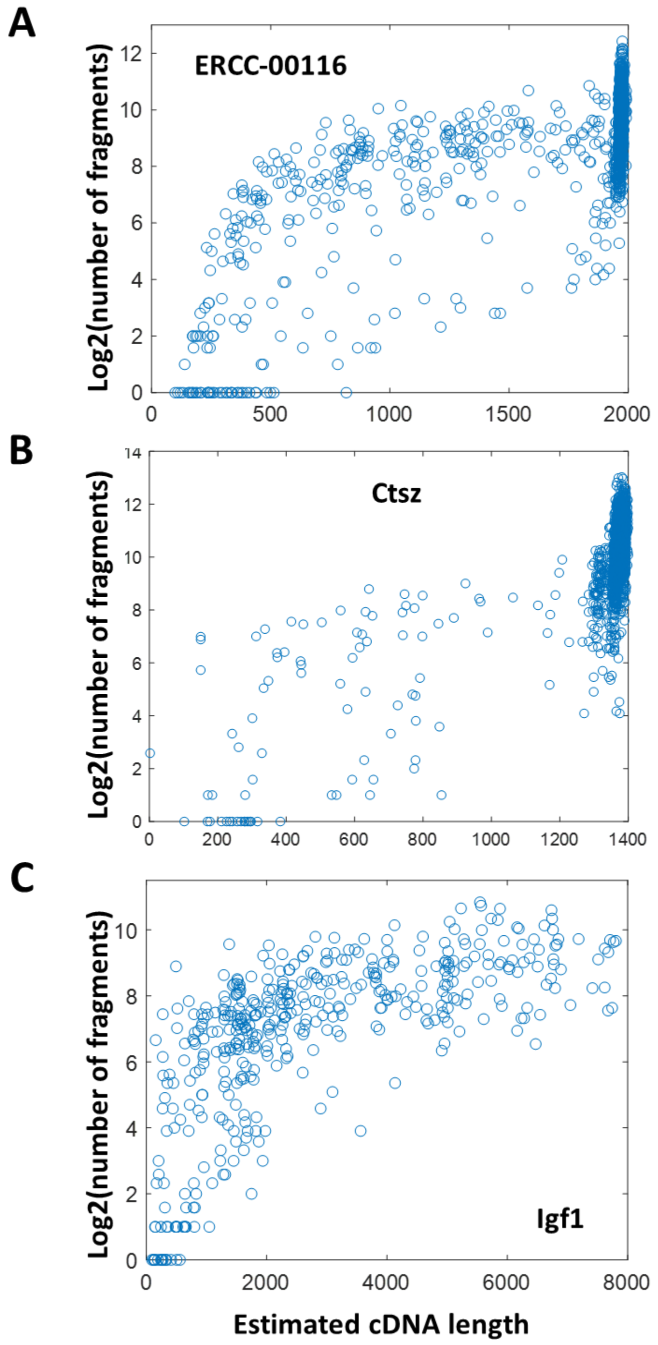


**Fig. S 3.** The number of a gene's sequenced fragments as a function of the estimated cDNA lengths. In the figure, one dot represents a fragment count of the gene in an individual cell.

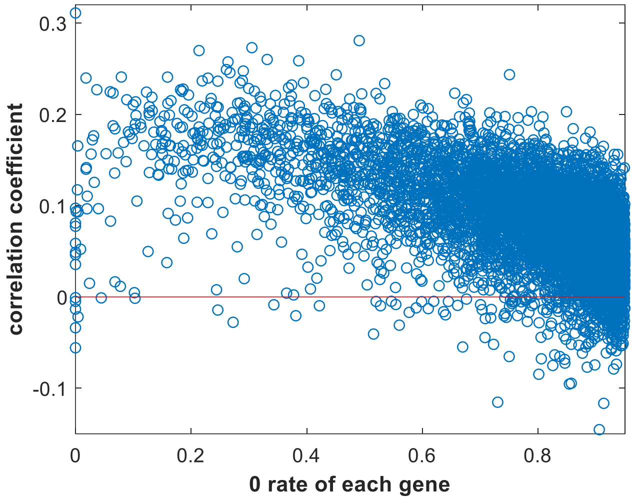


**Fig. S 4.** The correlation coefficient between sequencing depth (cell factor) and each gene's cDNA length (cDNA-length factor) across all cells. In the figure, one dot represents the correlation coefficient of a gene. The genes are ordered based on their 0 rates across cells. It shows that excepts for a small part of genes with high 0 rates (i.e., low expression rates), the corresponding cDNA lengths of most genes have a positive correlation with the cells' sequencing depths.

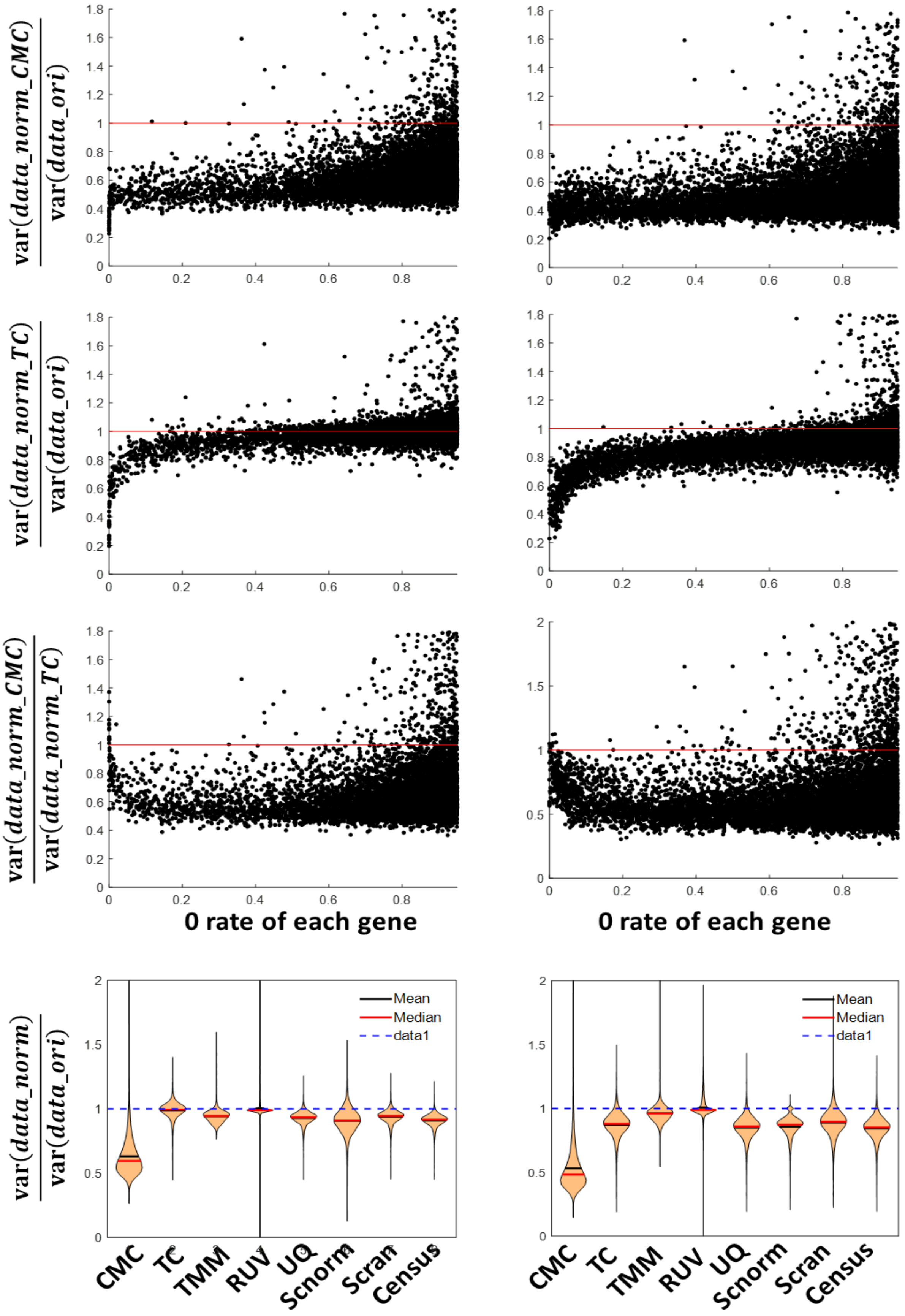
var(data_norm_CMC) / var(data_ori)
var(data_norm_TC) / var(data_ori)
var(data_norm_CMC) / var(data_norm_TC)
0 rate of each gene
0 rate of each gene
var(data_norm) / var(data_ori)
Mean
Median
data1
CMC
TC
TMM
RUV
UQ
Scnorm
Scran
Census

**Fig. S 5.** scRNA-seq data normalization for homogeneous microglia cells (first column) and homogeneous oligodendrocyte cells (second column). First two rows: variance change of each gene across all homogeneous cells before and after normalization (first row: CMC model; second row: Total Count (TC) method). Each dot represents the variance change of an individual gene. The y-axis is the ratio of variance after normalization to that of before normalization. The variances are all computed on the log2-transformed normalized data. The third row: comparison of each gene's variance after normalization by CMC model and by TC method, respectively. The last row: summary of all genes' variance changes after normalization via different methods.

| | #DEGs (FDR<0.05) | # overlapped with ground truth DEGs |
|---|---|---|
| **CMC** | 844 | 550 |
| **TC** | 259 | 197 |
| **TMM** | 318 | 220 |
| **RUV** | 219 | 162 |
| **UQ** | 493 | 354 |
| **Scnorm** | 599 | 408 |
| **Scran** | 536 | 372 |
| **Census** | 643 | 433 |

**Table S1.** Number of identified DEGs (under the threshold FDR<0.05) and their numbers of overlaps with the ground truth DEGs.

| | | | Gene 1 | Gene 2 | Gene 3 | ... |
|---|---|---|---|---|---|---|
| Dataset 1 | TF1 | Cell type1 | 0.5 | 0 | 0.8 | ... |
| Dataset 2 | TF1 | Cell type2 | 0.9 | 0.3 | 0 | ... |
| Dataset 3 | TF2 | Cell type2 | 0 | 0 | 0.6 | ... |
| Dataset 4 | TF3 | Cell type3 | 0.6 | 0 | 0 | ... |
| ... | ... | ... | ... | ... | ... | ... |

**Fig. S 6.** The TF-gene binding state tensor

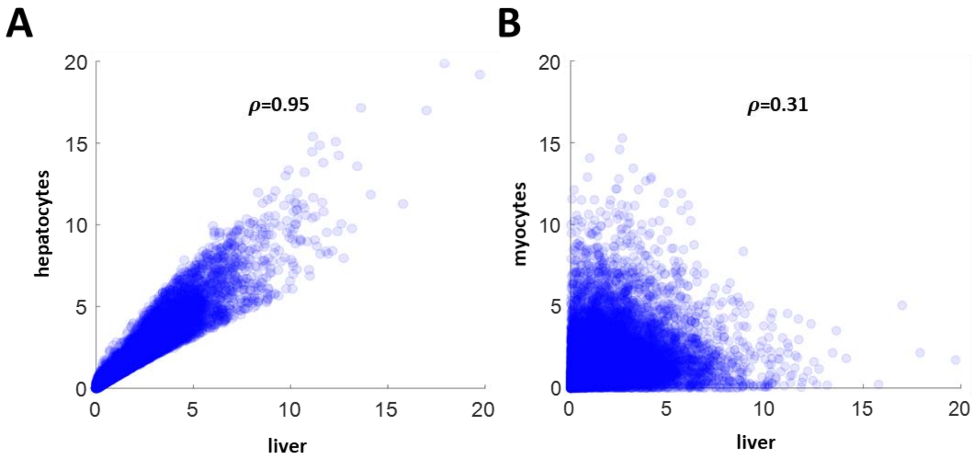


**Fig. S 7.** Comparison of the binding affinities of each gene in different tissues/cell types. (A) hepatocytes *vs.* liver. (B) myocytes *vs.* liver. In the figure, one dot represents the binding affinities of a gene in two different tissues/cell types.

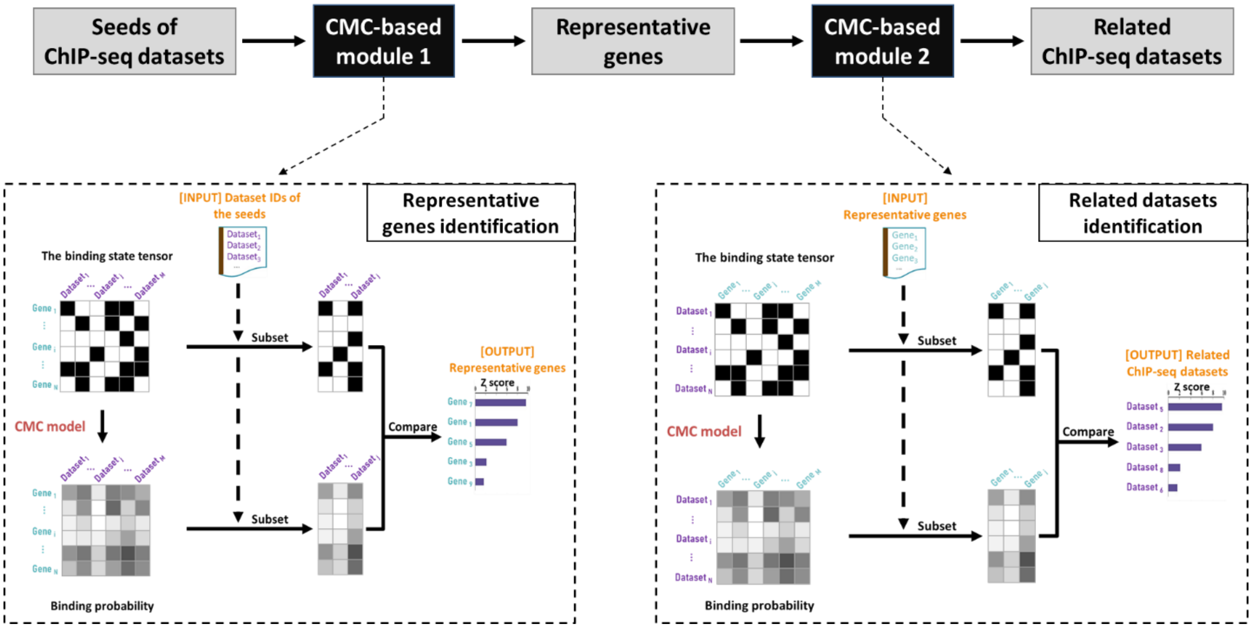


**Fig. S 8.** The CMC-based framework for related ChIP-seq dataset identification.

**CMC-based method**

| Rank | Dataset ID | p value | Cell Type of ChIP-seq datasets |
|---|---|---|---|
| 1 | EXP037772 | 6.34E-64 | Th0-cells |
| 2 | EXP038873 | 4.23E-63 | Th17-cells |
| 3 | EXP038870 | 5.54E-60 | Th17(beta)-cells |
| 4 | EXP038875 | 6.24E-59 | Th17-cells |
| 5 | EXP038596 | 1.63E-56 | Th17-cells |
| 6 | EXP037774 | 1.62E-55 | Th0-cells |
| 7 | EXP038868 | 1.02E-53 | Th17-cells |
| 8 | EXP038872 | 2.59E-53 | Th17-cells |
| 9 | EXP037769 | 2.81E-51 | Th17-cells |
| 10 | EXP030760 | 7.54E-50 | Th17-cells |
| 11 | EXP038866 | 3.49E-49 | Th0-cells |
| 12 | EXP038863 | 6.80E-49 | Th17-cells |
| 13 | EXP038597 | 4.74E-48 | Th17-cells |
| 14 | EXP031653 | 4.00E-46 | Th2-cells |
| 15 | EXP038888 | 3.54E-45 | Th0-cells |
| 16 | EXP038889 | 8.72E-45 | Th2-cells |
| 17 | EXP038864 | 1.23E-44 | Th17-cells |
| 18 | EXP038874 | 1.75E-44 | CD4+ T-cells |
| 19 | EXP038865 | 1.96E-44 | Th17-cells |
| 20 | EXP037659 | 5.96E-44 | Th0-cells |
| 21 | EXP038887 | 2.34E-43 | Th17-cells |
| 22 | EXP037664 | 2.75E-42 | Th17(beta)-cells |
| 23 | EXP038876 | 4.42E-42 | Th17-cells |
| 24 | EXP031638 | 9.31E-42 | Th17-cells |
| 25 | EXP038907 | 2.35E-41 | Th0-cells |

⋮

**Correlation-based method**

| Rank | Dataset ID | Correlation coefficient | p value | Cell Type of ChIP-seq datasets |
|---|---|---|---|---|
| 1 | EXP031718 | 0.512 | < 1E-300 | CD4+ T-cells |
| 2 | EXP034618 | 0.495 | < 1E-300 | acute myeloid leukemia |
| 3 | EXP037954 | 0.490 | < 1E-300 | bone marrow-derived macrophages |
| 4 | EXP053330 | 0.486 | < 1E-300 | MLL-AF9 Leukemic Cells |
| 5 | EXP034619 | 0.484 | < 1E-300 | acute myeloid leukemia |
| 6 | EXP033975 | 0.480 | < 1E-300 | thymocytes |
| 7 | EXP036591 | 0.479 | < 1E-300 | pre-B-cell progenitors |
| 8 | EXP038459 | 0.478 | < 1E-300 | CD4+ T-cells |
| 9 | EXP038748 | 0.473 | < 1E-300 | CD4+ T-cells |
| 10 | EXP033272 | 0.472 | < 1E-300 | mature megakaryocytes |
| 11 | EXP037290 | 0.471 | < 1E-300 | bone marrow-derived macrophages |
| 12 | EXP058860 | 0.469 | < 1E-300 | prostate cancer |
| 13 | EXP039859 | 0.463 | < 1E-300 | CH12.LX (Mouse lymphoma) |
| 14 | EXP033974 | 0.458 | < 1E-300 | thymocytes |
| 15 | EXP030589 | 0.456 | < 1E-300 | prostate |
| 16 | EXP037572 | 0.455 | < 1E-300 | Hoxb8-FL |
| 17 | EXP052117 | 0.454 | < 1E-300 | pro-T-tumor derived (Scid.adh.2c2 cells) |
| 18 | EXP058777 | 0.453 | < 1E-300 | bone marrow-derived macrophages |
| 19 | EXP031720 | 0.453 | < 1E-300 | CD4+ T-cells |
| 20 | EXP038886 | 0.452 | < 1E-300 | Th17-cells |
| 21 | EXP038956 | 0.451 | < 1E-300 | spleen B-cells |
| 22 | EXP031717 | 0.450 | < 1E-300 | CD4+ T-cells |
| 23 | EXP052102 | 0.449 | < 1E-300 | pro-T-tumor derived (Scid.adh.2c2 cells) |
| 24 | EXP037959 | 0.449 | < 1E-300 | bone marrow-derived macrophages |
| 25 | EXP037571 | 0.448 | < 1E-300 | Hoxb8-FL |

⋮

unrelated cell type

related cell type

**Fig. S 9.** Related ChIP-seq dataset identification results for regulatory T cells. Left: results of the proposed CMC-based method; Right: results of the correlation-based method.

**CMC-based method**

| Rank | Dataset ID | p value | Cell Type of ChIP-seq datasets |
|---|---|---|---|
| 1 | EXP038755 | 5.14E-116 | liver |
| 2 | EXP032039 | 1.03E-115 | liver |
| 3 | EXP031831 | 9.40E-114 | liver |
| 4 | EXP031823 | 2.31E-111 | liver |
| 5 | EXP031827 | 1.19E-104 | liver |
| 6 | EXP031835 | 1.32E-104 | liver |
| 7 | EXP031834 | 2.13E-101 | liver |
| 8 | EXP031992 | 8.87E-101 | liver |
| 9 | EXP031825 | 5.13E-100 | liver |
| 10 | EXP031828 | 2.35E-99 | liver |
| 11 | EXP031833 | 1.68E-98 | liver |
| 12 | EXP033796 | 4.32E-96 | liver |
| 13 | EXP031165 | 6.51E-96 | liver |
| 14 | EXP033740 | 1.26E-94 | liver |
| 15 | EXP033885 | 1.83E-93 | liver |
| 16 | EXP033792 | 2.18E-93 | liver |
| 17 | EXP031837 | 9.22E-91 | liver |
| 18 | EXP036075 | 9.23E-91 | liver |
| 19 | EXP033795 | 2.39E-90 | liver |
| 20 | EXP031824 | 8.87E-90 | liver |
| 21 | EXP033794 | 6.11E-89 | liver |
| 22 | EXP031830 | 1.04E-87 | liver |
| 23 | EXP031826 | 1.46E-86 | liver |
| 24 | EXP033600 | 1.80E-86 | liver |
| 25 | EXP035239 | 5.07E-86 | liver |

⋮

**Correlation-based method**

| Rank | Dataset ID | Correlation coefficient | p value | Cell Type of ChIP-seq datasets |
|---|---|---|---|---|
| 1 | EXP037737 | 0.687 | < 1E-300 | liver |
| 2 | EXP037733 | 0.683 | < 1E-300 | liver |
| 3 | EXP035238 | 0.643 | < 1E-300 | liver |
| 4 | EXP035239 | 0.642 | < 1E-300 | liver |
| 5 | EXP035240 | 0.638 | < 1E-300 | liver |
| 6 | EXP037419 | 0.629 | < 1E-300 | brown preadipocytes |
| 7 | EXP031994 | 0.621 | < 1E-300 | liver |
| 8 | EXP037418 | 0.614 | < 1E-300 | brown preadipocytes |
| 9 | EXP034594 | 0.612 | < 1E-300 | CD4+ T-cells |
| 10 | EXP038743 | 0.611 | < 1E-300 | bone marrow-derived macrophages |
| 11 | EXP037735 | 0.607 | < 1E-300 | liver |
| 12 | EXP031993 | 0.606 | < 1E-300 | liver |
| 13 | EXP031992 | 0.601 | < 1E-300 | liver |
| 14 | EXP033795 | 0.596 | < 1E-300 | liver |
| 15 | EXP033794 | 0.595 | < 1E-300 | liver |
| 16 | EXP033796 | 0.593 | < 1E-300 | liver |
| 17 | EXP037738 | 0.593 | < 1E-300 | liver |
| 18 | EXP038742 | 0.590 | < 1E-300 | bone marrow-derived macrophages |
| 19 | EXP035241 | 0.587 | < 1E-300 | liver |
| 20 | EXP038744 | 0.587 | < 1E-300 | bone marrow-derived macrophages |
| 21 | EXP038741 | 0.584 | < 1E-300 | bone marrow-derived macrophages |
| 22 | EXP037420 | 0.583 | < 1E-300 | brown preadipocytes |
| 23 | EXP038755 | 0.581 | < 1E-300 | liver |
| 24 | EXP030890 | 0.579 | < 1E-300 | 3T3-L1-derived adipocytes |
| 25 | EXP038740 | 0.576 | < 1E-300 | bone marrow-derived macrophages |

⋮

◁ related cell type

◁ unrelated cell type

**Fig. S 10.** Related ChIP-seq dataset identification results for hepatocytes. Left: results of the proposed CMC-based method; Right: results of the correlation-based method.

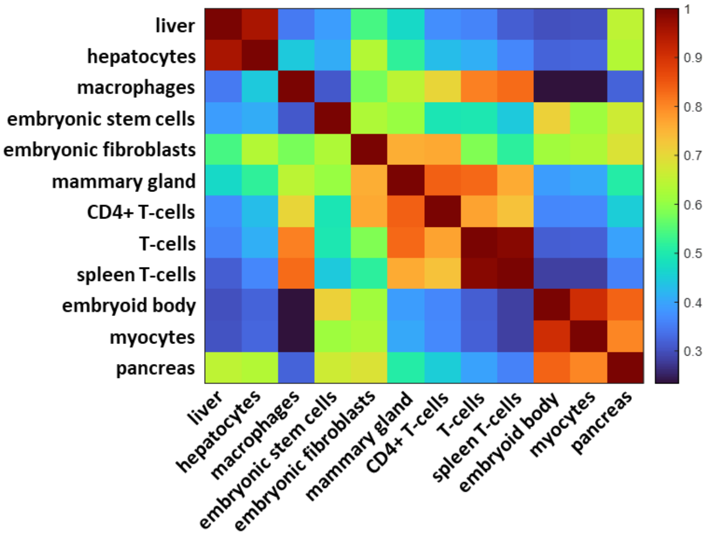


**Fig. S 11.** The correlation coefficient of the estimated genes' binding affinities between any two cell types. The correlation coefficients are shown as a heatmap.

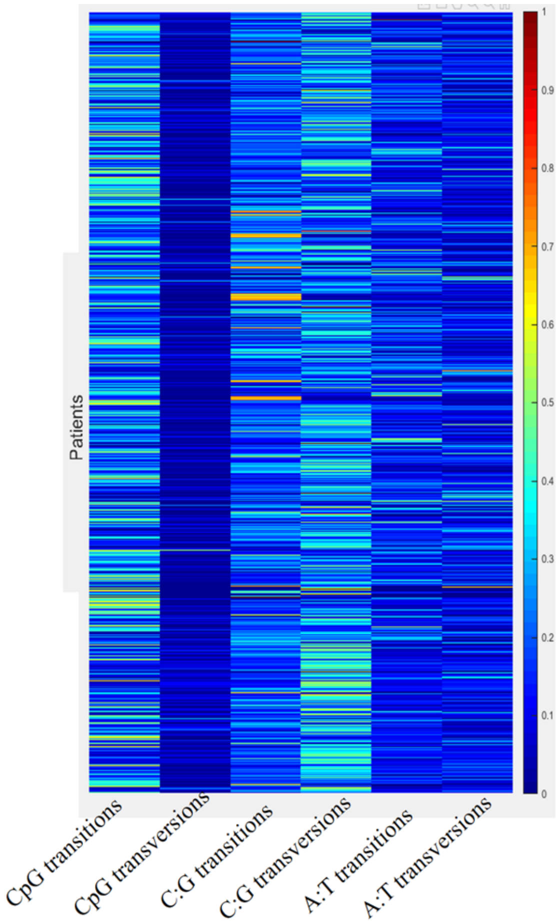


**Fig. S 12.** Heatmap of the mutation rate of each category within each patient. In this figure, the mutation rates are averaged across genes. In each patient, the mutation rates of the six categories are further normalized by dividing the sum of the six categories' mutation rates. The figure was plotted according to the mutation data of TCGA.

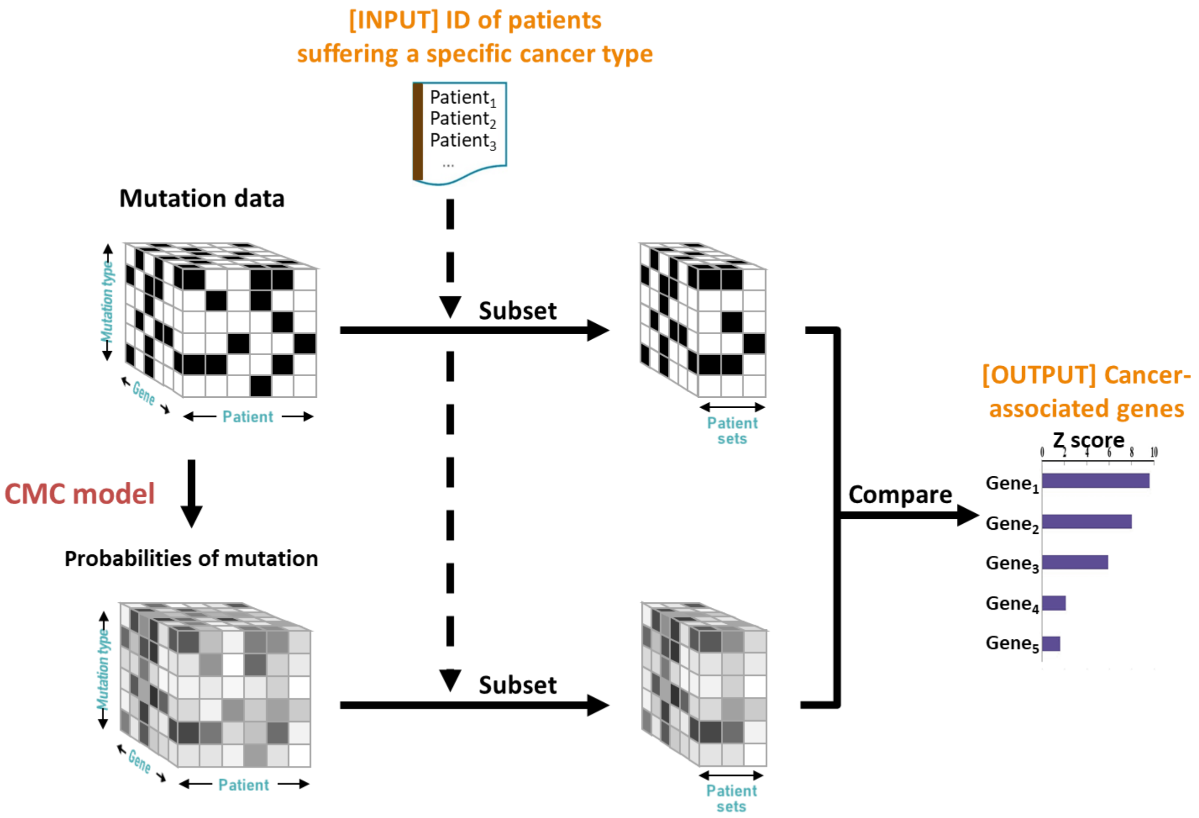


**Fig. S 13.** The framework of cancer-associated gene identification.

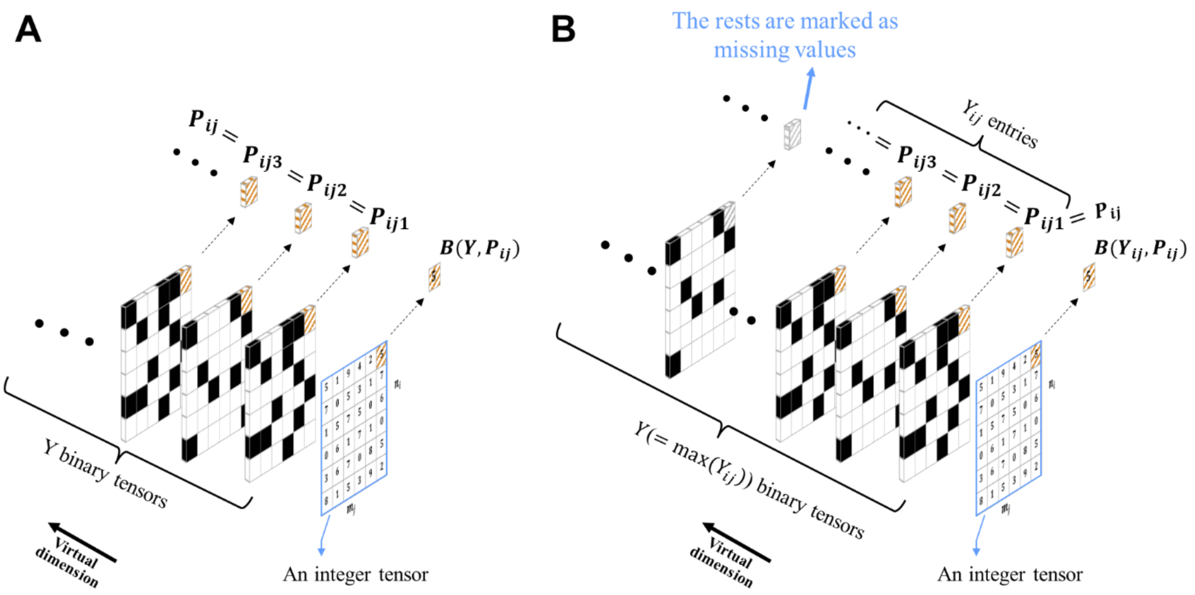


**Fig. S 14.** Illustration of decomposing a 2D integer tensor into a 3D binary tensor, in which a virtual dimension is introduced. Along with this virtual dimension, each entry in every individual is the results of a single trial in each entry. (A) The special case where the numbers of trials (*Y*) of each entry in the 2D integer tensor are the same. The length of the virtual dimension is equal to $Y$. Value of $\text{entry}_{ij}$ in the 2D integer tensor is the number of successes in a sequence of $Y$ independent Bernoulli trials (corresponding to $\text{entry}_{ij1}, \ldots, \text{entry}_{ijY}$ in the 3D binary tensor) with an identical success probability. (B) The general cases where $Y_{i,j}$s are different. The length of the virtual dimension is equal to $Y = \max(Y_{ij})$. For $\text{entry}_{ij}$ in the 2D integer tensor, among its corresponding entries ($\text{entry}_{ij1}, \ldots, \text{entry}_{ijY}$) in the 3D binary tensor, $Y - Y_{ij}$ of them ($\text{entry}_{ij\,(Y_{ij}+1)}, \ldots, \text{entry}_{ijY}$) are set as missing values.
